# The pro-tumoral and anti-tumoral roles of EphA4 on T regulatory cells and tumor associated macrophages during HNSCC tumor progression

**DOI:** 10.1101/2024.08.13.607778

**Authors:** Sophia Corbo, Diemmy Nguyen, Shilpa Bhatia, Laurel B. Darragh, Khalid N.M. Abdelazeem, Benjamin Van Court, Nicholas A. Olimpo, Jacob Gadwa, Justin Yu, Chloe Hodgson, Von Samedi, Enrique Sanz Garcia, Lillian Siu, Anthony Saviola, Lynn E. Heasley, Michael W. Knitz, Elena B. Pasquale, Sana D. Karam

## Abstract

Head and Neck Squamous Cell Carcinoma (HNSCC) is a deadly cancer with poor response to targeted therapy, largely driven by an immunosuppressive tumor microenvironment (TME). Here we examine the immune-modulatory role of the receptor tyrosine kinase EphA4 in HNSCC progression. Within the TME, EphA4 is primarily expressed on regulatory T cells (Tregs) and macrophages. In contrast ephrinB2, an activating ligand of EphA4, is expressed in tumor blood vessels. Using genetically engineered mouse models, we show that EphA4 expressed in Tregs promotes tumor growth, whereas EphA4 expressed in monocytes inhibits tumor growth. In contrast, ephrinB2 knockout in blood vessels reduces both intratumoral Tregs and macrophages. A novel specific EphA4 inhibitor, APY-d3-PEG4, reverses the accelerated tumor growth we had previously reported with EphB4 cancer cell knockout. EphA4 knockout in macrophages not only enhanced their differentiation into M2 macrophage but also increased Treg suppressive activity. APY-d3-PEG4 reversed the accelerated growth seen in the EphA4 knockout of monocytes but conferred no additional benefit when EphA4 was knocked out on Tregs. Underscoring an EphA4-mediated interplay between Tregs and macrophages, we found that knockout of EphA4 in Tregs not only decreases their activation but also reduces tumor infiltration of pro-tumoral M2 macrophages. These data identify Tregs as a primary target of APY-d3-PEG4 and suggest a role for Tregs in regulating macrophage conversion. These data also support the possible anti-cancer therapeutic value of bispecific peptides or antibodies capable of promoting EphA4 blockade in Tregs but not macrophages.

**Significance:** EphA4 in regulatory T cells has a pro-tumoral effect while EphA4 in macrophages plays an anti-tumoral role underscoring the necessity of developing biologically rational therapeutics.

## Introduction

The Eph gene family is the largest family of receptor tyrosine kinases (RTKs) and their corresponding ephrin ligands^1^. The Eph-ephrins are categorized into two subtypes, the receptors EphAs and EphBs, and their ligands ephrinAs and ephrinBs.^1^ In general, EphA receptors bind to ephrinA ligands, but there are exceptions to this. For example, EphA4 is known to bind to both ephrinA and ephrinB ligands, including ephrinB2.^2^ Both Eph receptors and their ligands are membrane bound, and both can signal through their cytoplasmic domain therefore capable of bidirectional signaling, with forward and reverse signaling occurring through the Eph receptor and ephrin ligand, respectively.^3,4^

Like many developmental genes, Eph-ephrin’s expression and function becomes mis-regulated in cancer development, progression, and metastasis.^5,6^ Their mis-regulation has been shown in a variety of cancers including but not limited to head and neck cancer, breast, lung, prostate, leukemia, and melanoma.^7^ In addition to promoting tumorigenesis some Eph-ephrins can act as tumor suppressors as well, highlighting the importance of understanding the biology of Eph-ephrin signaling during tumor progression.^7^ There have been several preclinical studies and clinical trials targeting Eph-ephrin signaling using small molecule inhibitors, synthetic peptides to inhibit Eph-ephrin binding, receptor tyrosine kinase inhibitors (RTKi), and monoclonal antibodies.^8^ Many of these therapies have shown an initial, but transient, period of response.^9–12^ Eph receptors have also been shown to contribute to therapeutic resistance.^6^ We previously developed an artificial model of resistance to targeted therapy where knockout/knockdown of EphB4 on the cancer cell displays an initial period of tumor growth delay but followed by accelerated tumor growth.^13^ As the Eph receptors share similar molecular structure with RTKs, they are the target of many RTKi^14^ making the understanding of their functional inhibition imperative to advancing cancer therapeutics. In addition to the cancer cell, Eph-ephrins are expressed on a variety of immune cells including monocytes, macrophages, dendritic cells, B cells, and T cells.^15^

Within the HNSCC tumor microenvironment (TME), it has been demonstrated that both regulatory T cells (Tregs) and macrophages, two immune populations with high EphA4 expression, play a key role in mediating tumor progression and resistance to immunotherapies^16–22^. As we are the first to characterize EphA4’s role within the TME, we unexpectedly observe opposing roles of EphA4 within these two immune compartments. Using genetically engineered mouse models, we determined that EphA4 knockout in Tregs decreased tumor volume and increased anti-tumoral immune response while EphA4 knockout on monocytes increased tumor volume, predominance, and pro-tumoral response. Finally, we utilized the newly developed specific peptide inhibitor, APY-d3-PEG4, as a novel translational therapeutic to target EphA4 and leverage its antitumoral-immune effects in HNSCC.

## RESULTS

### Oral cavity squamous cell carcinoma (OCSCC) patients display elevated EphA4 expression in Tregs

Given the observed an increase in EphA4 expression within the tumor microenvironment (TME) in our preclinical model of treatment resistance,^13^ we investigated the role of EphA4 in mediating such resistance. We first examined if expression of EphA4 and ligand binding partner eprhinB2^2^ correlated with survival of head and neck squamous cell carcinoma (HNSCC) patients. Survival analysis using publicly available data sets shows both EphA4 and ephrinB2 expression inversely correlate with survival in HNSCC patients (**Figure 1A**; p = 0.0082, HR = 1.69; 95% CI = 1.14 - 2.50). This suggested that EphA4 is a potential targetable receptor mediating progression in HNSCC. To further investigate the role of EphA4 during disease progression, we examined the effect of pharmacological inhibition of EphA4 with Sitravatinib in oral cavity squamous cell carcinoma (OCSCC) patients (PMID: 34599023).^23^ Sitravatinib is a tyrosine kinase inhibitor that inhibits a variety of receptor tyrosine kinases including EphA4.^14^ We performed mass cytometry (CyTOF) analysis on peripheral blood mononuclear cells (PBMCs) on 6 patients, from pre Sitravatinib treatment (Pre) and post Sitravatinib treatment (D15). CyTOF analysis revealed that at both Pre and D15, CD4+ T cells are the dominant lymphocyte within circulation (**Supplemental Figure 1A**). Further analysis of PBMCs revealed that CD4+ T cells display the highest EphA4 expression (**Figure 1B**). When examining the various subsets of CD4+ T cells, we observed pro-tumoral CD4+FoxP3+ and CD4+CD25+FoxP3+ regulatory T cells (Tregs) have increased EphA4 expression compared to anti-tumoral CD4+ T cells subsets (CD4+Tbet+ and CD4+IFNγ+) (**Figure 1C**).

**Figure 1.**
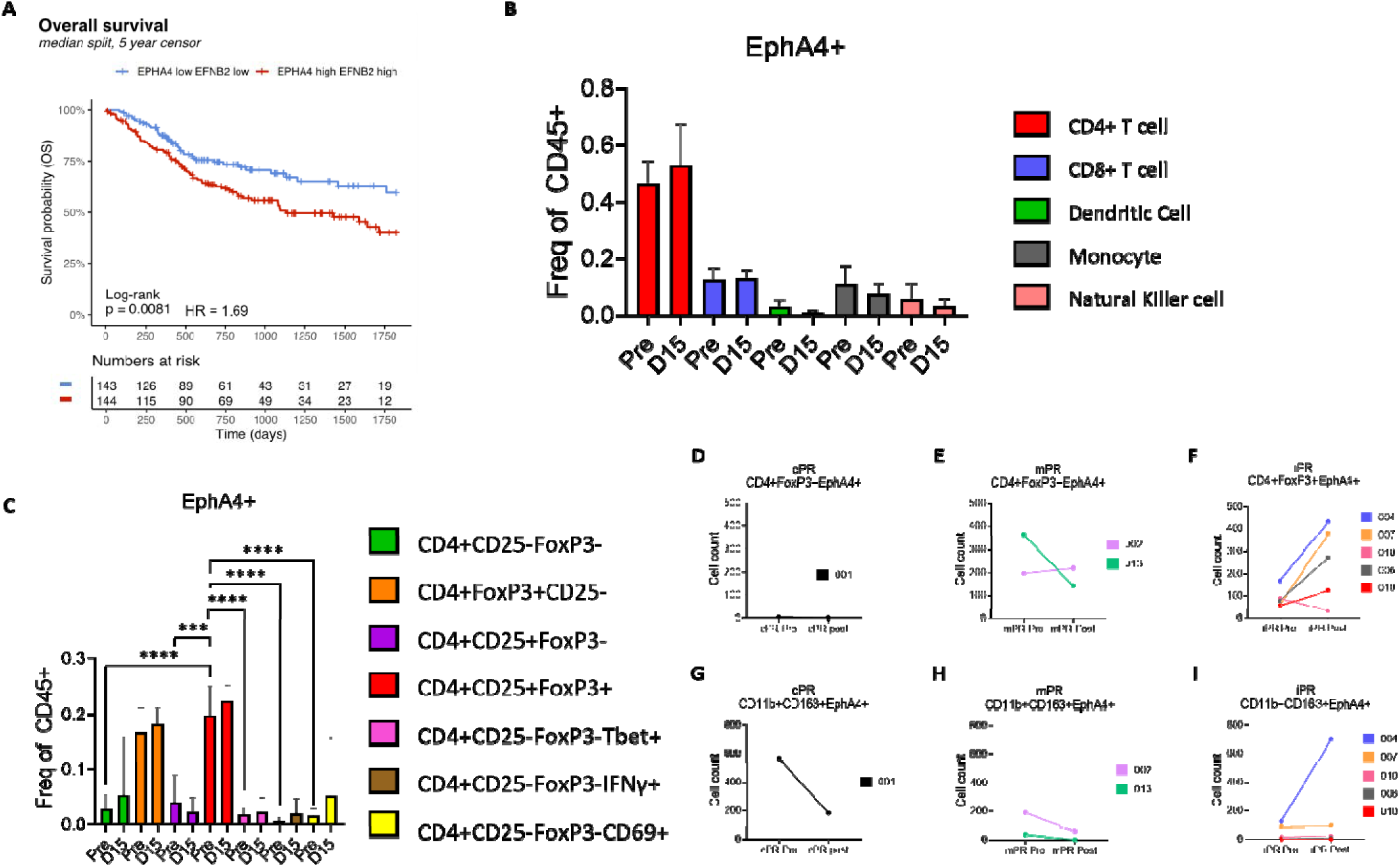
Oral cavity squamous cell carcinoma (OCSCC) iPR patients EphA4+ Tregs and pro-tumoral macrophages post Sitravatinib. **A)** TCGA survival probability on HNSCC patients examining expression of EphA4 and ephrinB2. Patients with high expression of EphA4 and eprhinB2 display a decrease survival probability. **B)** CyTOF was performed on PBMCs. EphA4 expression in various leukocytes from pretreatment (Pre) and 15 days post Sitravatinib treatment (D15) time points. Dendritic cells defined as CD45+CD3-D11c+, monocytes defined as CD45+CD3-CD14+, and natural killer cells defined as CD45+CD3-CD56+. **C)** CyTOF analysis of EphA4 expression on various subpopulations of CD4+ T cells. **D-I)** VECTRA Polaris imaging was performed on tumor tissue pre Sitravatinib treatment (pre) and post Sitravatinib treatment (post). Patients were categorized as either complete pathological response (cPR), major pathological response (mPR), or incomplete pathological response (iPR).

We next questioned which cell types within the tumor microenvironment expressed EphA4. To answer this question, we performed VECTRA Polaris multiplex immunofluorescence imaging of tumor tissue. We observed an increase in the frequency of EphA4+ Tregs in incomplete pathological response (iPR) patients, while complete pathological response (cPR) and major pathological response (mPR) displayed either a decrease or no change in EphA4+ Tregs (**Figure 1 D-F**). This was consistent with data from our published Phase I/Ib trial with neoadjuvant Durvalumab and radiation in oral cavity patients,^20^ showing a significant correlation between EphA4+Tregs and response to treatment (**Supplemental Figure 1B**). In addition to Tregs, we also observed EphA4 expression co-localize with pro-tumoral macrophages with cPR and mPR patients displaying a decrease in EphA4+ pro-tumoral macrophages and iPR patients displaying either an increase or no change in EphA4+ pro-tumoral macrophages (**Figure 1 G-I**). Due to the co-localization and trends with response to Sitravatinib, we focused our study on the role of EphA4 on Tregs and macrophages.

### Pharmacological inhibition of EphA4 inhibits the growth of EphB4 knockdown tumors

Taking into account that our preclinical treatment resistance model of HNSCC displays increased EphA4 mRNA and protein expression as well as EphA4 tyrosine phosphorylation^13^, and that EphA4 expression in Tregs and pro-tumoral macrophages increased for OCSCC iPR patients (**Figure 1**), we speculated that EphA4 promotes treatment resistance. Supporting this speculation is our previous work with the kinase inhibitors Dasatinib and Nilotinib, both reduced tumor growth in our model of treatment resistant HNSCC.^13^ However, Dasatinib^24^ and Nilotinib^25^ are ATP-competitive inhibitors that target many tyrosine kinases and are therefore not specific for EphA4. To investigate the effects of specific EphA4 pharmacological inhibition, we treated Moc2 EphB4 shRNA tumors and control shRNA tumors with APY-d3-PEG4 (**Figure 2A**), a novel PEGylated peptide developed to inhibit EphA4 signaling *in vivo*^26^. APY-d3-PEG4 is an EphA4 antagonist that targets with nanomolar binding affinity and high specificity the ephrin-binding pocket in the extracellular ligand-binding domain of EphA4^27^. The APY-d3 peptide moiety is highly resistant to plasma proteases and the conjugated polyethylene glycol (PEG) moiety confers prolonged half-life in the blood circulation.^27^ ^26,28^

**Figure 2.**
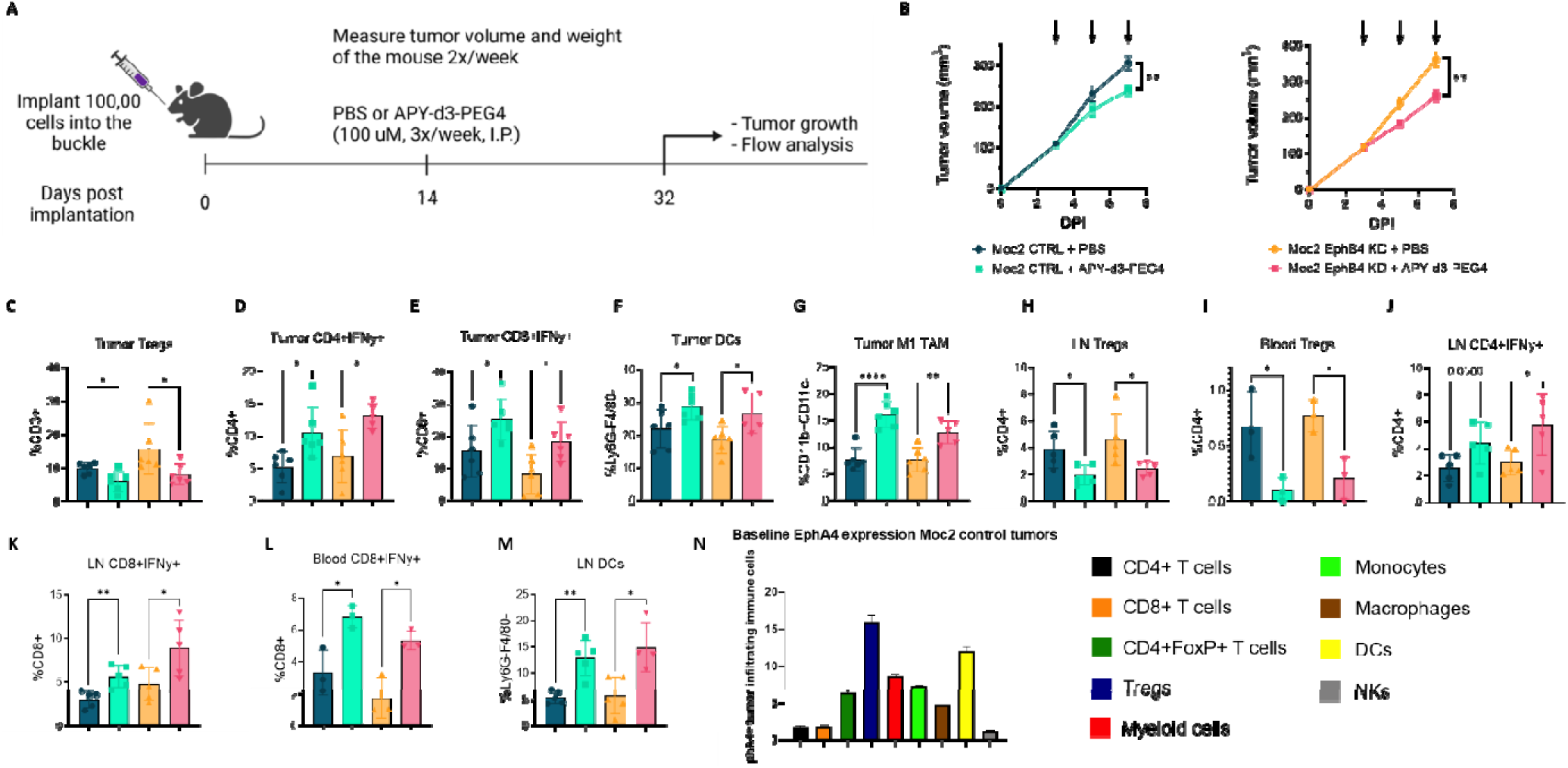
Pharmacological inhibition of EphA4 inhibits the growth of EphB4 knockdown Moc2 tumors and increases pro-inflammatory effects within the TME. **A)** Moc2 control and Moc2 EphB4 knockdown cells were implanted in the buccal region of C57BL/6 mice followed by treatment of PBS or 250 μL 100 μM APY-d3-PEG4 treatment three times per week by intraperitoneal injection. **B)** Tumor growth curve, caliper measurements were used to measure tumor growth. Significance was determined using a parametric two-tailed T test. **C-M)** Flow cytometry was performed on day 11 post implantation at the time of sacrifice. Green represents Moc2 control mice treated with PBS control, purple represents Moc2 control knockout tumors treated with APY-d3-PEG4, blue represents Moc2 EphB4 knockdown tumors treated with PBS control, and red represents Moc2 EphB4 knockdown tumors treated with APY-d3-PEG4. **C)** Definition of Tregs as follows CD45+CD3+CD4+CD25+FoxP3+. **F)** Definition of dendritic cells as follows CD45+CD3-Ly6G-F4/80-D11c+. **G)** Definition of M1 TAM as follows CD45+CD3-CD11b+CD11c-iNOS+. **N)** Flow cytometry analysis on Moc2 control shRNA tumors. CD4+ T cells (CD45+CD3+CD4+), CD8+ T cells (CD45+CD3+CD8+), Tregs (CD45+CD3+CD25+FoxP3+), Myeloid cells (CD45+Gr1+), monocytes (CD45+CD11b+Ly6C+), macrophages (CD45+CD11b+F4/80+), dendritic cells (CD45+CD11c+), and natural killer cells (NKs) (CD45+NKp46+). For statistical analysis p values are as follows, * = p ≤ 0.05, ** = p ≤ 0.01, *** = p ≤ 0.001, and **** = p ≤ 0.0001.

Building on our previous work^13^, we hypothesized that EphA4 pharmacological inhibition will only be effective at reducing accelerated tumor growth when EphA4 expression increases on the cancer cell. To examine the effects of pharmacological inhibition of EphA4 in EphB4 knockdown Moc2 tumors, we administered 250 μL of 100 μM APY-d3-PEG4 intraperitoneally three times per week (**Figure 2A**). A significant reduction in tumor growth was observed in both control and EphB4 knockdown tumors (**Figure 2B, Supplemental Figure 2 A-D**), suggesting that the efficacy of the peptide is independent of the level of cancer cell EphA4 expression. To further examine the efficacy of APY-d3-PEG4 on tumor growth, we used a second model, the MEER HNSCC cell line, where EphA4 expression does not increase with EphB4 knockdown (**Supplemental Figure 3 A-C**). Following the same dosing regimen used in Figure 1A (**Supplemental Figure 2E**), we found that treatment of MEER EphB4 knockout and MEER control knockout tumors with APY-d3-PEG4 does reduce tumor growth (**Supplemental Figure 2 F-J**). These data suggest that the pharmacological efficacy of EphA4 inhibition is independent of level of expression on the cancer cell and implicate TME involvement for target cells.

### The novel peptide APY-D3-PEG reduces cancer cell growth associated with EphB4 knockout and increases anti-tumoral immune effects within the TME

Given that EphA4 is expressed in various leukocytes^29–32^, we explored the effects of this novel APY-d3-PEG peptide on the immune TME of EphB4 knockdown and control tumors (**Figure 2 C-M, Supplemental Figure 4,5**). Flow cytometric analysis revealed a reduction in pro-tumoral Treg frequency in both control and EphB4 knockdown tumors (**Figure 2C**). This reduction was accompanied by an increase in antitumoral CD4+ T helper 1 (Th1) cells (**Figure 2D**) suggesting a potential role of EphA4 in mediating CD4+ T cell polarization between Th1 and Treg phenotype. Treatment with APY-d3-PEG4 peptide was also associated with increased CD8+ effector T cell function (**Figure 2E**) and increased dendritic cells and M1 macrophages (**Figure 2F, G**). These data suggest that EphA4 is a likely culprit in promoting accelerated tumor growth by promoting a pro-tumoral immune response, most likely by enhancing Treg suppression. In addition to the TME, we also examined the tumor draining lymph node and blood compartments. In concordance with the TME, we observed a reduction in Treg frequency accompanied by an increase in Th1 CD4+ T cells (**Figure 2 H-J**), with associated increase in CD8+ effector T cell function **(Figure 2K, L**), and increased frequency of dendritic cells (**Figure 2M**). In accordance with these changes and confirming target immune populations of APY-d3-PEG4, flow cytometric analysis showed high levels of EphA4 protein on Tregs, monocytic cell populations including monocytes and macrophages, and dendritic cells (**Figure 2N**). Collectively, these data advance the hypothesis that pharmacological inhibition of EphA4 reduces Tregs-mediated immunosuppression and allows for antigen presenting cells to promote effector T cell function and exert anti-tumoral immune effects.

### Inhibition of EphA4 on Tregs limits tumor progression, decreases Treg activation and proliferation, facilitating a TME with CD4 helper phenotype and anti-tumoral macrophages

Given the high level of expression of EphA4 on Tregs (**Figure 2N**), and their published role in mediating tumor growth and resistance to therapy in several head and neck models^16–18,33^, we hypothesized that EphA4 is promoting Treg immunosuppression. To investigate the role of EphA4 on Tregs during tumor progression, we utilized a flox/cre genetically engineered mouse model (GEMM) to knockout (KO) EphA4 on FoxP3 expressing Tregs (EphA4^fl/fl^FoxP3^Cre^). Moc2 EphB4 knockdown and Moc2 control knockdown cells implanted into EphA4^fl/fl^FoxP3^Cre^ and EphA4^WT/WT^FoxP3^WT^ mice were compared for effect on tumor growth effects and immune TME (**Figure 3A**). We observed a significant decrease in tumor volume for both Moc2 EphB4 knockdown and control knockdown tumors during EphA4 knockout on Tregs (**Figure 3B, Supplemental Figure 6 A-D**). This suggests that EphA4 could mediate accelerated tumor growth by modulating Treg function.

**Figure 3.**
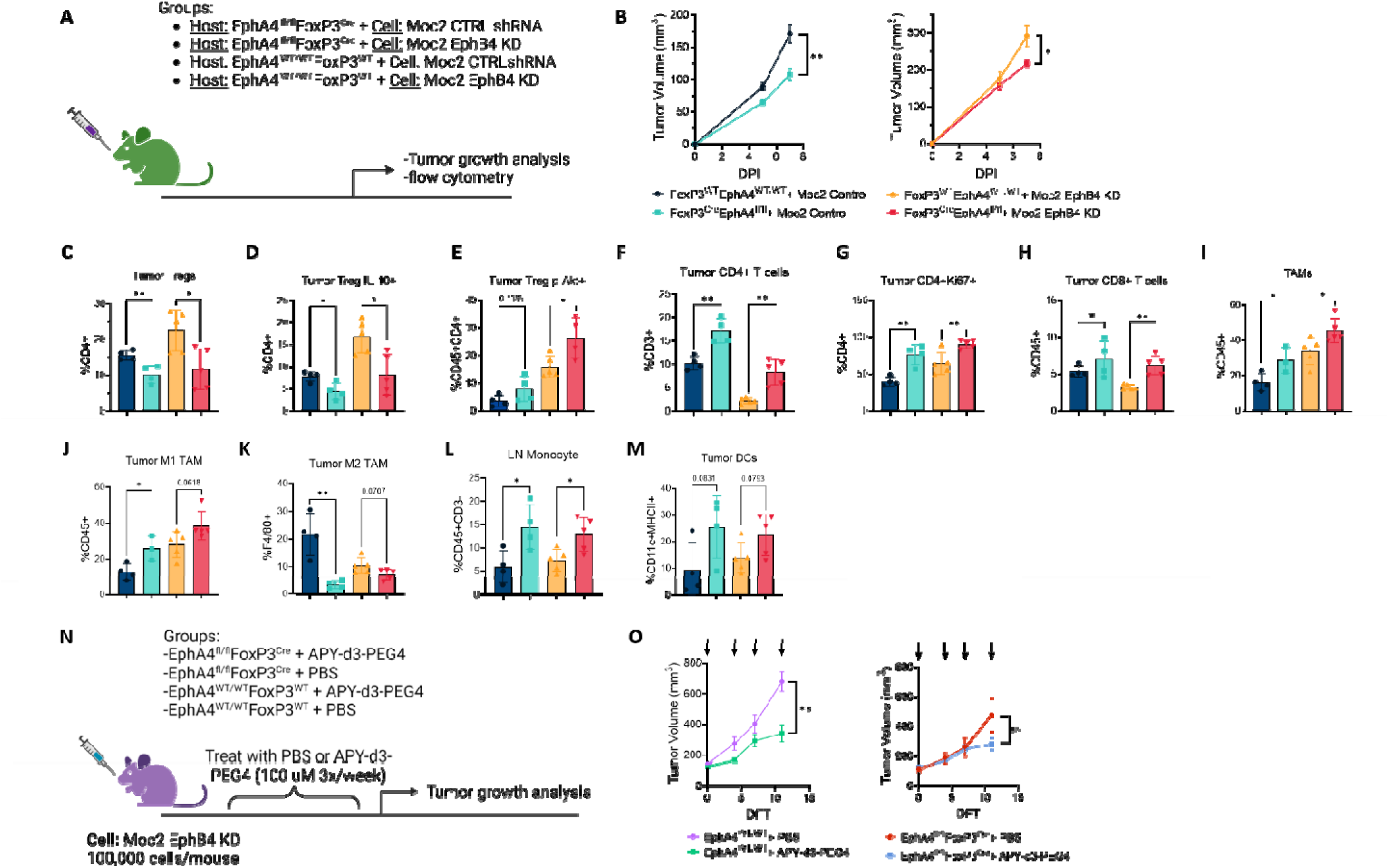
Inhibition Of EphA4 on Tregs limits tumor progression and Treg immunosuppression. **A)** Moc2 control and Moc2 EphB4 knockdown cells were implanted in the buccal region of EphA4^fl/fl^FoxP3^Cre^ and EphA4^WT/WT^FoxP3^WT^ mice. **B)** Tumor growth curve, both Moc2 control and Moc2 EphB4 knockdown tumors displayed decreased tumor growth when EphA4 is knockout on Tregs. **C-M)** Flow cytometry was performed on tumors and tumor draining lymph nodes on day 9 post implantation. **C)** Definition of Tregs as follows CD45+CD4+CD25+FoxP3+. **E)** Definition of Treg phosphorylated-Akt as follows CD45+CD4+CD25+p-AKT+. **I)** Definition of tumor associated macrophages as follows CD45+F4/80+. **J)** Definition of M1 TAM as follows CD45+F4/80+iNOS+. **K)** Definition of M2 tumor associated macrophage as follows CD45+F4/80+CD163+. **L)** Definition of a lymph node macrophage as follows CD45+Ly6C+Ly6G-. **M)** Definition of dendritic cells as follows CD45+Ly6G-CD11c+MHCII+CD103+. **N)** Moc2 EphB4 shRNA cells were implanted in the buccal region of EphA4^fl/fl^FoxP3^Cre^ and EphA4^WT/WT^FoxP3^WT^ mice followed by treatment of PBS or 100 uM APY-d3-PEG4 treatment three times per week. **O)** Tumor growth, significance was determined using an unpaired parametric T test. For statistical analysis p values are as follows, * = p ≤ 0.05, ** = p ≤ 0.01, *** = p ≤ 0.001, and **** = p ≤ 0.0001.

We next explored the effects of EphA4 KO on Tregs on the immune TME of EphB4 knockdown and control tumors (**Figure 3 C-M, Supplemental Figure 7, 8**). Within the TME during EphA4 loss on Tregs, we observed decreased Treg frequency and IL-10 production (**Figure 3 C, D**). In addition, we observed increased phosphorylated Akt (p-Akt) on CD4+CD25+ T cells (**Figure 3E**). It has been previously shown that activation of Akt signaling in Tregs represses the translation of key Treg associated genes.^34^ These data suggest a fragile Treg^34^ phenotype during EphA4 loss. Correlating with decreased Treg function, we observed an increase in CD4+ T cell frequency and proliferation (**Figure 3 F, G**). In addition to CD4+ T cells, we observed a significant increase in the frequency of CD8+ T cells during EphA4 knockout on Tregs in Moc2 EphB4 knockout tumors (**Figure 3H**). A notable increase in M1 macrophages in both the TME and tumor draining lymph node was also observed along with decreased M2 macrophages intratumorally (**Figure 3 I-L**). Antigen presenting dendritic cells (MHCII+CD103+CD11c+) also showed an increased trend intratumorally during EphA4 loss on Tregs (**Figure 3M**). Together, these data suggest that selectively targeting EphA4 in Tregs decreases immunosuppression leading to enhanced antigen presentation and effector T cell function.

We next questioned if the therapeutic benefit observed during APY-d3-PEG4 treatment is mediated by EphA4 on Tregs. To examine if pharmacological inhibition of EphA4 in Tregs is mediating decreased tumor growth, we treated EphA4^fl/fl^FoxP3^Cre^ and EphA4^WT/WT^FoxP3^WT^ mice with APY-d3-PEG4 (**Figure 3N**). We found that genetic knockout of EphA4 on Tregs abolished APY-d3-PEG4’s therapeutic benefit (**Figure 3O, Supplemental Figure 6 E-H**). This indicates the role of EphA4 in Tregs is critical in mediating accelerated tumor growth during EphB4 loss on the cancer cell.

### EphA4 knockout Tregs display decreased Treg function

To examine how EphA4 knockout in Tregs leads to decreased Treg frequency within the tumor microenvironment (**Figure 3C**), we performed proteomic analysis on tumor Tregs containing EphA4 knockout. We harvested Tregs from the Moc2 EphB4 knockdown tumors and tumor draining lymph nodes from EphA4^fl/fl^FoxP3^Cre^ and DEREG mice. After isolating the Treg population using fluorescence-activated cell sorting (FACs) we subjected murine Tregs to proteomic analysis (**Figure 4A**). Our proteomic analysis revealed a decrease in heat shock proteins (HSPs) Hsp90aa1, Hsp90ab1, and Hspa8 (**Figure 4 A-D**). Various HSPs have been implicated in promoting Treg immunosuppression by inducing Treg production of IL-10.^35^ A decrease in HSPs with EphA4 knockout on Tregs suggests that these Tregs have decreased immunosuppression compared to when EphA4 expression is maintained.

**Figure 4.**
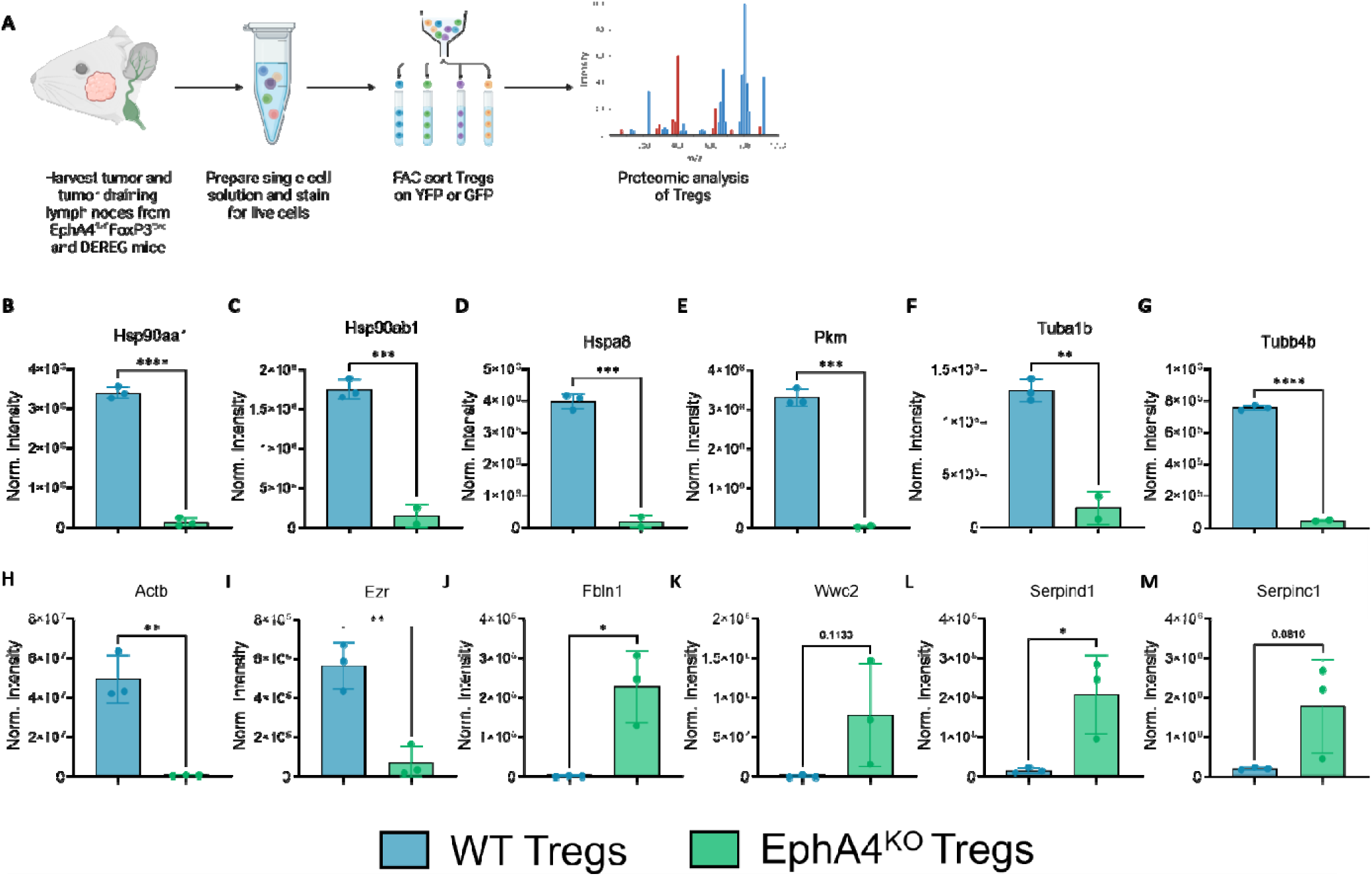
EphA4 knockout Tregs display decreased Treg function. **A)** Schematic of Treg isolation for proteomics. Tregs were harvested from Moc2 EphB4 knockdown tumors and tumor draining lymph nodes. Single cell solutions were then subjected to FACs to isolate Tregs. **B-H).** Proteomic analysis of knockout Tregs and wild type Tregs. EphA4

We also observed a decrease in pyruvate kinase (PKM), an enzyme that converts ADP to ATP in glycolysis,^36^ with EphA4 knockout (**Figure 4E**). This suggests Tregs with EphA4 knockout have decreased glycolysis, which Tregs utilize to rapidly produce ATP needed for Treg migration.^37,38^ In support of decreased PKM with EphA4 knockout we observed decreased cytoskeletal function marked by decreased Tubulin alpha-1B chain (Tuba1b), Tubulin beta-4B chain (Tubb4b), Actinβ (Actb), and Ezrin (Ezr) (**Figure 4 F-I**).

Furthermore, our proteomic analysis revealed an increase in Fibulin-1 (Fbln1) and Protein WWC2 (Wwc2) (**Figure 4J, K**). Both Fbln1 and Wwc2 are negative regulators of Treg immunosuppression mediated by NOTCH^39^ and Hippo^40,41^ pathways, respectively. In agreement with EphA4 KO correlation with decreased function EphA4 KO Tregs display increased Serpind1 and Serpinc1 (**Figure 4L, M**). Serpins are known to inhibit granzyme release,^42^ Tregs utilize granzyme to inhibit effector T cells.^43^ All together these data suggest, EphA4 in Tregs correlates with enhanced Treg immunosuppression.

### Genetic knockout of EphA4 on monocytes increases tumor progression, immunosuppressive macrophages, and promotes Tregs expansion within the TME

Given the high level of EphA4 in macrophages (**Figure 2N**), we sought to determine if EphA4 loss in macrophages would contribute to decreased tumor growth. We utilized a flox/cre GEMM to genetically knockout EphA4 in LysM expressing cells (EphA4^fl/fl^LysM^Cre^) and implanted Moc2 EphB4 knockdown and Moc2 control knockdown cells into the buccal mucosa of EphA4^fl/fl^LysM^Cre^ and EphA4^WT/WT^LysM^WT^ mice for tumor growth effect and flow cytometric analysis (**Figure 5A**). We unexpectedly found that when EphA4 is genetically knockout in macrophages, a significant increase in tumor growth was observed relative to control for both Moc2 EphB4 knockdown and Moc2 control knockdown tumors (**Figure 5B, Supplemental Figure 9 A-D**). These findings were validated in a second model, with MEER EphB4 knockout implanted in EphA4^fl/fl^LysM^Cre^ mice (**Figure 5 C, D, Supplemental Figure 9E, F**). We hypothesized that the accelerated tumor growth with EphA4 inhibition in macrophages is driven by reprogrammed intra-tumoral macrophage phenotype, whereby inhibition of EphA4 is reprogramming TME monocyte differentiation towards a pro-tumoral macrophage phenotype.

**Figure 5.**
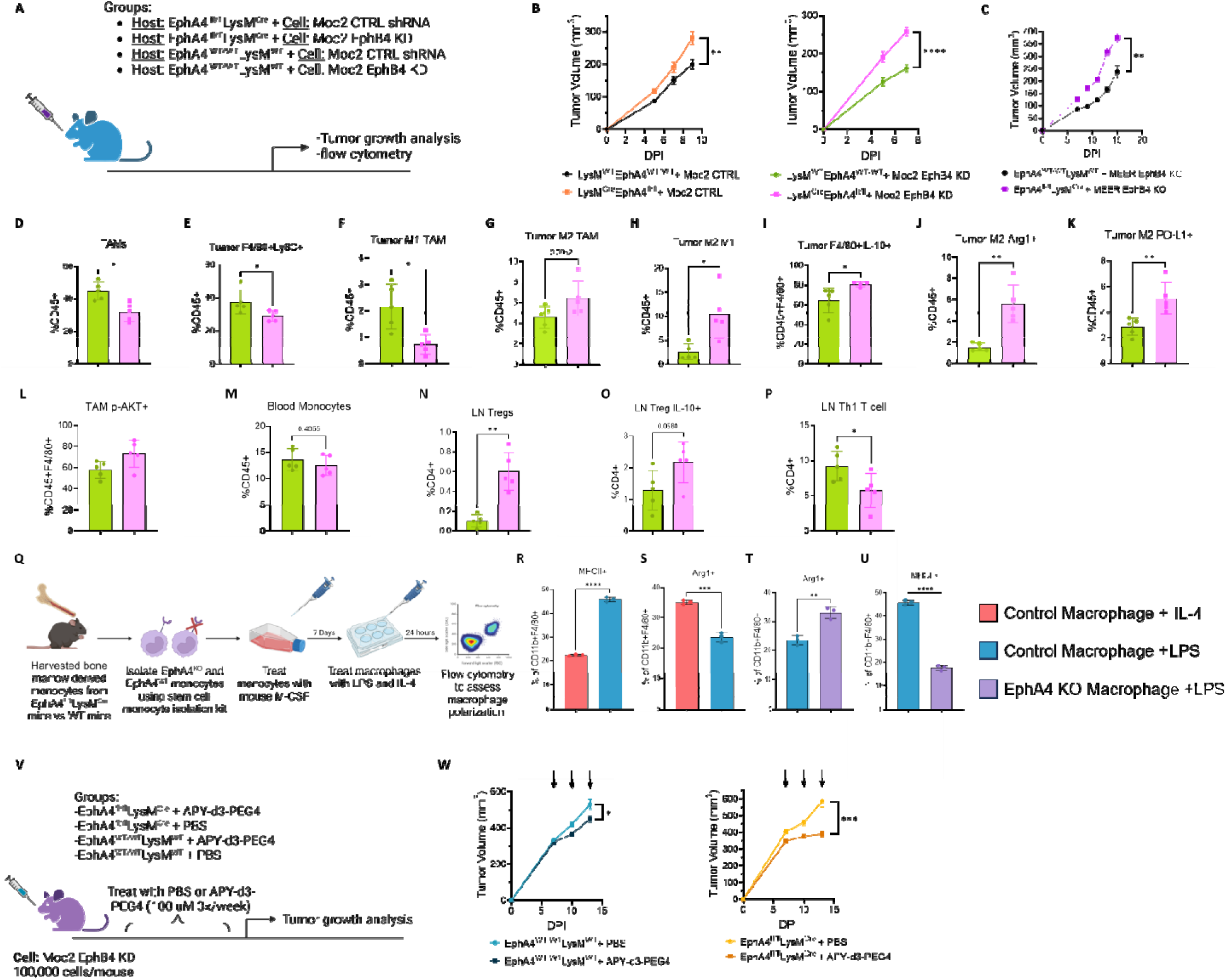
Genetic knockout of EphA4 on monocytes increases tumor progression, immunosuppressive macrophages, and promotes Tregs expansion within the TME. **A)** Moc2 control shRNA or Moc2 EphB4 shRNA cells were implanted into the buccal mucosa of EphA4^WT/WT^LysM^WT^ or EphA4^fl/fl^LysM^Cre^ mice. **B)** Tumor growth, during EphA4 knockout on LysM+ monocytes we observe accelerated tumor growth for both Moc2 control and Moc2 EphB4 knockdown tumors. Significance was determined using Mann-Whitney T test. **C)** Tumor growth, tumors with EphA4 knockout monocytes display reduced tumor volume. **D-P)** Flow cytometry was performed on tumors on day 9 post implantation. **D)** Definition of tumor associated macrophages as follows CD45+F4/80+. **F)** Definition of M1 tumor associated macrophages as follows CD45+F4/80+Ly6G-Ly6C+IFNγ+. **G)** Definition of M2 tumor associated macrophages as follows CD45+F4/80+CD1 3+. **M)** Definition of monocytes as follows CD45+Ly6C+Ly6G-. **P)** Definition of Th1 T cell as follows CD4+Tbet+IFNγ+. **Q)** Schematic for bone marrow derived macrophage *in vitro* polarization assay. **R-U)** Quantification of macrophage polarization using flow cytometry. **V)** Moc2 EphB4 shRNA cells were implanted in the buccal region of EphA4^WT/WT^LysM^WT^ or EphA4^fl/fl^LysM^Cre^ mice followed by treatment of PBS or 100 µM APY-d3-PEG4 treatment three times per week. **W)** Tumor growth, APY-d3-PEG4 treatment showed reduced tumor volume for EphA4^fl/fl^LysM^Cre^ mice. Significance was determined using an unpaired T test. For statistical analysis p values are as follows, * = p ≤ 0.05, ** = p ≤ 0.01, *** = p ≤ 0.001, and **** = p ≤ 0.0001.

To test this hypothesis, we performed multicompartmental flow cytometric analysis on tumor, tumor draining lymph nodes, and blood (**Figure 5 D-P, Supplemental Figure 10, 11**). Our analysis showed that intratumorally, a significant decrease in overall tumor associated macrophages (TAMs) was observed when EphA4 is knockout on LysM+ cells (**Figure 5D, E**). Notably, a significant decrease in anti-tumoral M1 macrophages and an increase in the frequency of pro-tumoral M2 macrophages were observed yielding a significantly increased M2/M1 macrophage ratio (**Figure 5 F-I**). The activation of M2 macrophages was also observed as evidenced by increased IL-10, Arg1, and PD-L1 when EphA4 is genetically knockout on macrophages compared to control (**Figure 5 J-K**). Interestingly, we observe the same pattern of increased p-Akt during EphA4 knockout on macrophages as we observed during EphA4 inhibition on Tregs (**Figure 5L**), suggesting a potential role where EphA4 inhibits Akt activation. In macrophages, Akt activation has been associated with driving expression of various M2 related genes.^44^ Together these data suggest that EphA4 expression promotes the polarization of macrophages towards a pro-tumoral M2 phenotype. To confirm that EphA4 is mediating macrophage polarization and not M2 recruitment to the TME we also examined the frequency of circulating monocytes. We observed no change in the frequency of circulating monocytes (**Figure 5M**), suggesting that EphA4 is mediating macrophage differentiation or polarization. Within the tumor draining lymph node, we observed increased Tregs and Treg IL-10 production (**Figure 3 N, O**). In addition, we observed decreased Th1 cells during EphA4 knockout on monocytes (**Figure 5P**). These data confirm that genetic deletion of EphA4 on macrophages promotes a pro-tumoral phenotype leading to increased tumor growth. Our data also support macrophage to Treg cross talk within the TME via IL-10 signaling, as macrophage secreted IL-10 has been demonstrated to promote Tregs expansion.^45^

We next examined if EphA4 expression on monocytes mediates polarization between anti-tumoral M1 and pro-tumoral M2 macrophages. To test this, we harvested bone marrow derived monocytes from EphA4^WT/WT^LysM^Cre^ and EphA4^fl/fl^LysM^Cre^ mice, differentiated monocytes into macrophages, and stimulated them with LPS and IL-4 to polarize macrophages into M1 and M2, respectively (**Figure 5Q**). We found when control macrophages were stimulated with LPS there was an M1 phenotype (**Figure 5R**) compared to stimulation with IL-4, which resulted in an M2 phenotype (**Figure 5S**). In contrast, when EphA4 KO macrophages were stimulated with LPS, these macrophages remained in an M2 phenotype and were unable to polarize to an M1 phenotype (**Figure 5T, U**). These data suggest that EphA4 knockout on macrophages promotes a pro-tumoral M2 phenotype.

We next tested whether this increase in pro-tumoral macrophages observed with EphA4 knockout on macrophages is mitigating the effects of APY-d3-PEG4 (**Figure 2B**). To test this, we treated EphA4^fl/fl^LysM^Cre^ and EphA4^WT/WT^LysM^WT^ mice with APY-d3-PEG4 (**Figure 5V**). We observed decreased tumor growth in EphA4^fl/fl^LysM^Cre^ mice treated with APY-d3-PEG4 compared to vehicle treated mice (**Figure 5W, Supplemental Figure 9 G-J**). While these data clearly demonstrate that EphA4 exerts an anti-tumoral M1 effect on macrophages, they also show that the primary target APY-d3-PEG4 are the Tregs as evident by the reversal of accelerated tumor growth effects seen in the EphA4^fl/fl^LysM^Cre^. These data also underscore the regulatory effects Treg inhibition has on macrophage polarization, with loss of EphA4 favoring M2-like polarization, but when Tregs are targeted with APY-d3-PEG4, it negates this polarization effect. The findings from **Figure 5W** along with the inability of APY-d3-PEG4 to inhibit accelerated tumor growth promoted by EphB4 loss in EphA4^fl/fl^Foxp3^Cre^ mice (**Figure 3O**) suggest the anti-tumoral effects of APY-d3-PEG4 are mediated by inhibition of EphA4 on Tregs.

### EphA4: ephrinB2 signaling promotes macrophage polarization

To examine how EphA4 interacts with its ligand, ephrinB2, which is predominantly expressed on the vessel, we utilized the EFNB2^fl/fl^Tie2^Cre^ GEMM (**Figure 6A**).^13^ Using flow cytometric analysis of tumor revealed that there was no change in TAM frequency but there is was a significant increase in TAM iNOS+ upon ephrinB2 knockout on VECs (**Figure 6B, C, Supplemental Figure 13A**). This suggests that ephrinB2 plays a role in TAM polarization within the TME. We also observed a significant decrease in circulating Ly6C^High^Ly6G-monocytes during eprhinB2 knockout on VECs (**Figure 6D, Supplemental Figure 13B**). Ly6C^High^Ly6G-monocytes typically develop into proinflammatory macrophages when they migrate into tissues.^46^ The decrease in proinflammatory circulating monocytes could suggest increased migration of this population into the TME. In agreement with this hypothesis, we observed increased proinflammatory TAMs within the TME upon ephrinB2 knockout on VECs (**Figure 6D**). There was no difference in circulating Ly6C^Low^Ly6G-monocytes during eprhinB2 knockout on VECs (**Figure 6E, Supplemental Figure 13B**). These data suggest that ephrinB2 signaling could promote monocyte polarization and recruitment to the TME.

**Figure 6.**
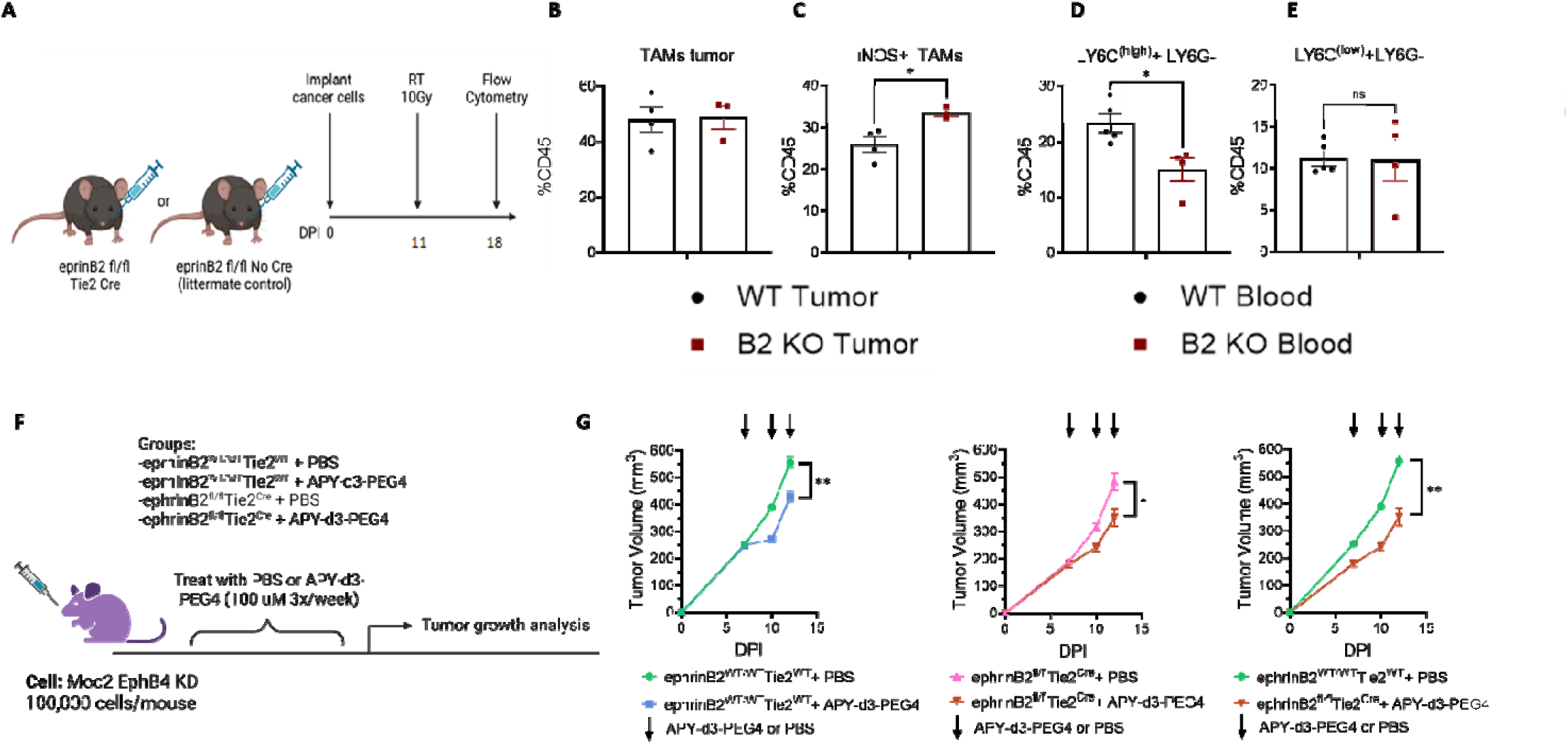
EphA4:EFNB2 signaling promotes accelerated tumor growth. **A)** Experimental design for studying the TME of EFNB2^fl/fl^Tie2^Cre^ mice. **B)** Quantification of tumor associated macrophages (TAMs) by CD45^+^CD11b^+^F4/80^+^. **C)** Quantification of M1 macrophages in the TME, defined by iNOS expression (TAMs: CD45^+^CD11b^+^F4/80^+^). **D-E)** Quantification of LY6C^(high)^ and LY6C^(low)^ monocytes in the blood. **F)** Moc2 EphB4 knockdown cells were implanted in the buccal region of ephrinB2^fl/fl^Tie2^Cre^ and littermate control mice followed by treatment of PBS or 100 µM APY-d3-PEG4 treatment three times per week by intraperitoneal injection. **G)** Tumor growth, significance was determined using an unpaired parametric T test. For statistical analysis p values.

### Vascular knockout of ephrinB2 synergizes with EphA4 pharmacological inhibition

With data indicating that pharmacological inhibition of EphA4 (**Figure 2B**) and/or genetic knockout of ephrinB2 on VECs^13^ leads to reduced tumor volume, we next investigated if additional benefit could be obtained with dual inhibition of both EphA4 and ephrinB2. To accomplish this dual inhibition, Moc2 EphB4 knockout cells were implanted into EFNB2^fl/fl^Tie2^Cre^ mice^13^ and pharmacologically inhibited EphA4 using APY-d3-PEG4 (**Figure 6F**). Dual inhibition of ephrinB2 and EphA4 led to greater decrease in tumor volume compared to single inhibition of either ephrinB2 or EphA4 (**Figure 6G, Supplemental Figure 12 A-D**). These data suggest a potential role of EphA4 receptor interacting with ephrinB2 ligand to mediate accelerated tumor growth, and dual inhibition of this receptor-ligand interaction could provide a targetable axis for recurrent HNSCC.

## DISCUSSION

With head and neck squamous cell carcinoma (HNSCC) displaying a heterogenous tumor microenvironment (TME) with several cell types and overlapping gene expression profiles,^47^ pleiotropic effects driving tumor growth can be the result of genes having opposing effects on different cell types or similar effects on cells with opposing functions. We have previously shown this redundancy in the TME of HNSCC when loss of EphB4 on cancer cells promotes accelerated tumor growth coinciding with increased expression of EphA4.^13^ We observe a reversal of the accelerated tumor growth phenotype when EphA4 is pharmacologically inhibited (**Figure 2B**). Consistent with previous work,^13^ we also found that eprhinB2, a high affinity binding ligand to both EphB4 and EphA4,^2^ is expressed on endothelial cells and its knockout on these cells decreases tumor growth.^13^ Given that EphA4 is a binding partner to ephrinB2, and its upregulation with EphB4 knockout/knockdown, we sought to determine the role EphA4 plays during HNSCC progression, hypothesizing that its upregulation serves a compensatory role for disease progression. To our knowledge, this is the first report to describe EphA4’s role in mediating tumor immune response and to dissect its action, *in vivo*, by cellular compartment. Our unexpected findings highlight the cell specific, and often opposing, effects this gene plays on tumor immune microenvironment and its sequential effect on tumor growth.

Within the TME, EphA4 was highly expressed in Tregs and macrophages. While EphA4 knockout on Tregs, either genetically or pharmacologically, led to decreased Treg suppressive activation and enhanced CD4+ T cell anti-tumoral immunity, targeting EphA4 in macrophages established the reverse. Not only do these data emphasize the cellular specific effects of EphA4’s function, but they also highlight the important interplay between the adaptive and innate immune system. Our data demonstrate that EphA4 knockout in Tregs controls macrophage polarization rendering them anti-tumoral. Similarly, knockout of EphA4 in macrophages not only rendered them pro-tumoral, but it also increased Tregs infiltration and suppressive function. While it could be argued that these results demonstrate cooperative and bi-directional effects of macrophage-Treg interaction, our data with pharmacological inhibition of EphA4 in either the EphA4^fl/^flLysM^Cre^ or the EphA4^fl/fl^FoxpP3^Cre^ mice demonstrate the far-reaching role Tregs play in regulating immune response. Specifically, the APY-d3-PEG4 peptide failed to add benefit when EphA4 is knocked out in Tregs (EphA4^fl/fl^FoxpP3^Cre^), but it reversed the effect of accelerated tumor growth in the LysM^Cre^EphA4^fl/fl^ mice. Similarly, when EphA4 was knocked out in EphA4^fl/fl^LysM^Cre^ mice, increased Tregs activation was noted accompanying the increase in pro-tumoral tumor associated macrophages within the TME.

In support of our tumor growth and flow cytometry findings in EphA4^fl/fl^FoxpP3^Cre^ mice, was our proteomic analysis of EphA4 knockout Tregs. One important finding from our analysis was the decreased Pkm with EphA4 knockout, suggesting a decrease in glycolysis. In Tregs glycolysis is tightly mediated by CD28 activation of the PI3K-Akt pathway.^37^ The PI3K-Akt promotes the activation of glucokinase, which further interacts with actin to mediated cytoskeletal rearrangements.^37^ These cytoskeletal rearrangements are critical for Treg migration. Therefore, we purpose inhibition of EphA4 on Tregs decreases glycolysis, further leading to decreased Treg immunosuppression in target tissues due to the inability to migrate. In addition to decreased migratory ability due to decreased glycolysis, we observed a decrease in key players that mediate cytoskeletal rearrangements. These cytoskeletal rearrangements are critical for Treg migration. For Tregs to effectively promote immunosuppression appropriate Treg positioning is required, therefore Treg migration is critical.^48^ Here we show that EphA4 knockout Tregs have decreased cytoskeletal function, therefore limiting their migratory potential. With limited migration Tregs these are unable to migrate within the tumor microenvironment to promote immunosuppression.

Tregs have long been appreciated as key mediators of tumor recurrence and treatment resistance by promoting immunosuppression within the TME.^16–19,49^ Heavily explored has been their role in blunting antigen presenting cells (APCs) function^16,50^ therefore, impairing their ability to present antigen to T cells thus limiting T effector cell (Teff) activation.^16–19,51^ While our findings show that EphA4 in Tregs limits APC activation, our results underscore the underexplored interplay between Tregs and macrophages. Macrophages can be broadly classified into two—the proinflammatory M1 macrophages which arise in the presence of cytokines like IFN-γ, and the anti-inflammatory CD163 positive M2 macrophages, which secrete immunosuppressive signals such as IL-10 and TGF-β.^45^ Previous studies have shown that in co-culture experiments, Tregs reduce M1 cytokines by monocytes while increasing M2 cytokines.^52^

Literature examining EphA4’s role in Tregs’ development and function is scant. In patients with Type I diabetes, EphA4 expression on Tregs has been shown to be reduced compared to conventional CD4 Tregs.^53^ Analysis of placental bed uterine Tregs (uTregs) showed lower levels of EphA4 expression compared to those from malignant breast or colon cancer patients.^54^ In human Tregs with CRISPR-mediated gene knock-in, EphA4 expression was also noted to be reduced in expression.^55^ As Tregs exists in different states, notably a memory Treg and activated or effector Treg state,^56–58^ these expression data showing reduced EphA4 in autoimmune states suggest that EphA4 likely upregulated on effector T cell states. Although no data exist in the cancer literature, our findings are consistent with EphA4 correlating with an activated Treg phenotype with increased CD44, CD25, and IL10 expression. IL10 secretion by Tregs has been shown to serve as a critical modulator of innate immune cell function.^59^ This has been extensively studied in mediating dendritic cell anergy but has also been hypothesized to polarize macrophages towards a pro-tumoral phenotype.^59^

While the effect of EphA4 on macrophage polarization can be an indirect one through enhanced Tregs activation,^59^ our data also support a direct mechanism of action. We show that knockout of EphA4 in macrophages resulted in marked increase in pro-tumoral macrophages and significant reduction of anti-tumoral macrophages. Consistent with these findings are results from a neural inflammation brain injury model where in bone marrow chimeras lacking EphA4 showed polarization of macrophages towards an anti-inflammatory-M2-like macrophages.^31^ In atherogenesis and wound healing models, it has also been shown that EphA4 promotes monocyte adhesion and inflammation by interacting with its ligands on the endothelium to promote their adhesion.^32,60^ Concordant with these findings, inhibiting ephrinB2 a binding partner of EphA4 using EFNB2^fl/fl^Tie2^Cre^ mice showed decreased circulating Ly6C^hi^ monocytes, but with increased M1 macrophages within the TME (**Figure 6 D, E**). In contrast, EphA4^fl/fl^LysM^cre^ shows no change in total circulating monocytes (**Figure 5M**) but increased in M2 macrophages (**Figure 5G**). Furthermore, the addition of EphA4 blocking peptide in the background of ephrin B2 mice yielded reduced tumor growth. These data collectively suggest that while adhesion and trans-endothelial migration may play an important role in coordinating the fate of intra-tumoral macrophages especially as it pertains to vascular EFNB2, they also underscore the importance of EphA4 in directly dictating monocyte anti-tumoral phenotypic state.

The implications of this work are profound. The model used here is one of progression in the face of EphB4 knockout and as such recapitulates the compensatory responses that arise in the face of treatment resistance. Perhaps most pertinent though, our findings underscore how the simplicity of nature’s functional redundancy presents a challenge for therapeutic targeting, both temporally and spatially. Pan-targeting of a protein that is used to activate multiple cells, immunosuppressive and immunostimulatory immune cells, can result in opposing effects that can reduce or negate any therapeutic benefit from such targeting. This calls for next generation bispecific peptides or antibodies capable of isolating EphA4’s blockade to Tregs thus avoiding macrophages. Next-generation Treg-cell therapies tailored to knocking out EphA4 and delivery via CAR-Treg or adoptive cell therapy. Finally, our results extend beyond cancer therapies and hold promise for autoimmune and infectious disease, as it relates to both macrophages and Tregs.

## Methods/Materials

### Cell Culture

The HNSCC murine cell lines were obtained as follows: Moc2 cell line from Dr. Ravindra Uppaluri (Dana-Farber Cancer Institute, Boston, MA), Ly2 cell line from Dr. Nadarajah Vigneswaran (University of Texas Health Science Center, Houston, TX), and bEND.3 cells were obtained from the lab of Dr. Jordan Jacobelli (Anschutz Medical Campus, Aurora, CO). All cell lines in this study were within 12 passages and tested for mycoplasma contamination prior to their use in the experiments. Generation of Moc2 EphB4 knockdown and MEER EphB4 knockout cell lines were generated as described in.^13^

### Tumor inoculation of buccal mucosa

Cancer cell lines were implanted into the buccal mucosa as described in^61^. Moc2 cell lines were implanted at a concentration of 1 x10^5^ cells per mouse. MEER cell lines were implanted at a concentration of 2.25 x10^5^ cells per mouse.

### Animal Handling

All animals were handled according to IACUC protocol.

### Administration of APY-d3-PEG4

APY-d3-PEG4 was obtained from Biosynth. APY-d3-PEG4 stock (concentration 500 μM) was diluted to 100 μM in 250 μL in PBS. APY-d3-PEG4 was administered by intraperitoneal injection (IP) three times per week.

### Generation of genetically engineered mouse models

EphA4^fl/fl^ mice (Epha4^tm1.1Bzh^/J) (Strain #: 012916), FoxP3^Cre^ mice (B6.129(Cg)-Foxp3^tm4(YFP.iCre)Ayr^/J) (Strain #: 016959), and LysM^Cre^ mice (B6.129P2-Lyz2^tm1(Cre)Ifo^/J) (Strain #: 004781) all were purchased from The Jackson Laboratory. Starting at 5 weeks old EphA4^fl/fl^ mice were bred with FoxP3^Cre^ and LysM^Cre^ mice to generate EphA4^fl/fl^FoxP3^Cre^ and EphA4^fl/fl^LysM^Cre^ mice. Ear snips were used to validate genotype with Transnetyx Inc. Positive and negative controls were used to validate genotype results.

### Flow cytometry

Flow cytometry was performed as described in.^16^ For analysis of immune cells the following antibodies were used: Arg1 (R&D Systems Cat #: IC5868F) CD3 (BioLegend Cat #: 100218), CD4 (Invitrogen Cat #: 48-0041-82), CD11b (BioLegend Cat #: 101235), CD11c (BD Pharmingen Cat #: 550261), CD45 (Invitrogen Cat #: 56-0451-82), CD80 (BD OptiBuild Cat #: 746775), CD163 (BioLegend Cat #: 155316), F4/80 (BioLegend Cat #: 123118), FoxP3 (Invitrogen Cat #: 58-5773-82), IFNγ (BD Horizon Cat #: 612769), IL-10 (BioLegend Cat #: 505031), iNOS (BioLegend Cat #: 696806), Ki67 (Invitrogen Cat #: 15-5698-82), Ly6C (BioLegend Cat #: 128010), Ly6G (BioLegend Cat #: 127641), MHCII (BioLegend Cat #: 107624), Phosphorylated AKT (BD Phosflow Cat #: 562599), PD-L1 (BD Horizon Cat #: 752339), TNFα (Invitrogen Cat #: 505-7321-82), CD103 (BioLegend Cat #: 121435), CD8 (BD Horizon Cat #: 564422), Tbet (Invitrogen Cat #: 606-5825-82), CD25 (BD Horizon Cat #: 564023).

### Western blot

For western blotting cells were grown in a monolayer to prepare protein lysates. Lysates were run on gels, transferred to membranes, and probed with primary antibodies as described in.^62^ Anti-EphA4 (1:1000 cat # 37-1600) was purchased from Invitrogen. Anti-β-Actin HRP conjugated (1:5000 cat # 12262S) was purchased from Cell Signaling Technology.

### Cell line proteomics

Cells were grown in a monolayer, harvested, and washed four times with PBS. Samples were run on the single-cell mass spectrometer.

### Mass cytometry (CyTOF) on PBMCs

PBMCs were obtained from Dr. Lillian Siu. PBMCs were isolated according to.^23^ Staining of PBMCs for CyTOF was performed as previously described.^20^ Analysis of CyTOF data was done in FlowJo (v10.10.0).

### Treg Proteomics

Tregs were harvested from EphA4^fl/fl^FoxP3^Cre^ and DEREG mice containing Moc2 EphB4 knockdown tumors. Tumors and tumor draining lymph nodes were harvested from these mice for Treg isolation. Tumors and tumor draining lymph nodes were strained through a 70 µm strainer to make a single cell solution. Live cells were stained using LIVE/DEAD fixable aqua dead cell stain kit (REF #: L34966). Tregs were subjected to fluorescence-activated cell sorting (FACs), using the SONY MA900. Tregs were sorted out based on yellow florescent protein (YFP) for EphA4^fl/fl^FoxP3^Cre^ Tregs and green florescent protein (GFP) for DEREG Tregs. After FACs we performed mass spectrometry using the Brucker timsTOF single cell proteomics (SCP) mass spectrometer.

### Macrophage Polarization Assay

Harvest femurs from mice and remove all the tissue surrounding the bone. Take a 0.5 mL Eppendorf tube and poke a hole in the bottom of tube. Place this 0.5 mL Eppendorf in a 1.5 mL Eppendorf tube. Place femurs into 0.5 mL Eppendorf and close lid. Centrifuge at 4000 RCF for 15 seconds. Resuspend bone marrow samples in RPMI and proceed to EasySep Mouse Monocyte Isolation Kit (from StemCell Technologies Cat #: 19861A). Add 10 mL of RPMI-C 10% + 10 ng/mL Macrophage Colony Stimulating Factor (M-CSF) to T-75 flask. Plate 1 x 10^6^ cells per T-75 flask. Put the plates in a 37 °C incubator (day 0). Macrophages were plated, cultured, harvested from plates, and stimulated with IL-4 and LPS as previously described.^63^

## Supporting information

All supplemental figures will be used for the link to the file on the preprint site

## Acknowledgements

We would also like to acknowledge our funding sources: R01DE028529, R01CA28465, R01DE028529, 1P50CA261605-01 (SDK), F31 DE029997 (LBD), T32DC012280 (SC).

---

**Supplemental Figure 1.**
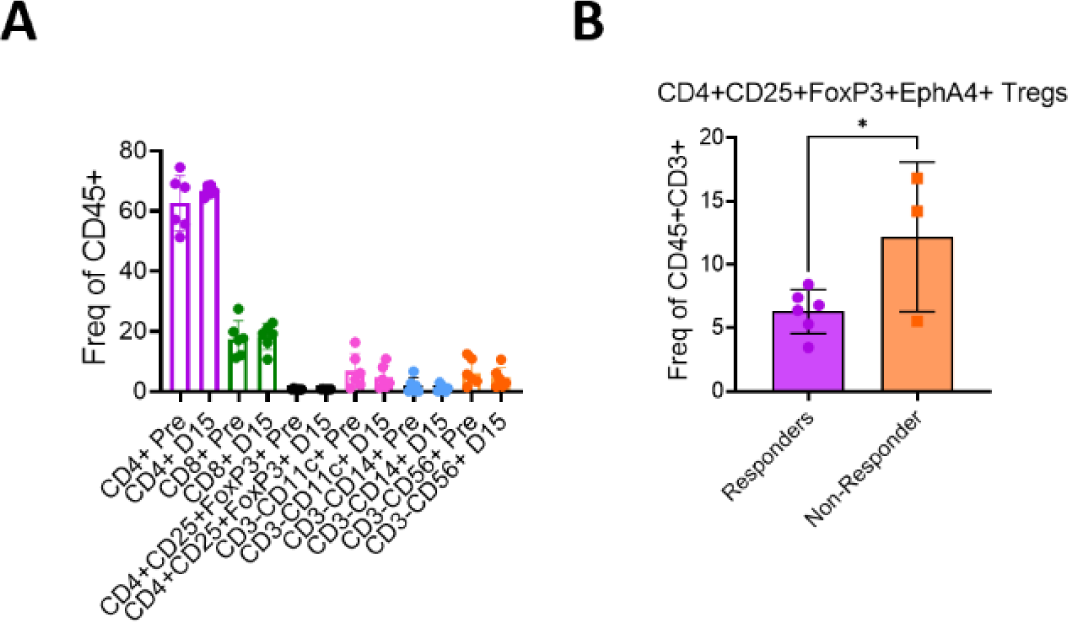

**Supplemental Figure 2.**
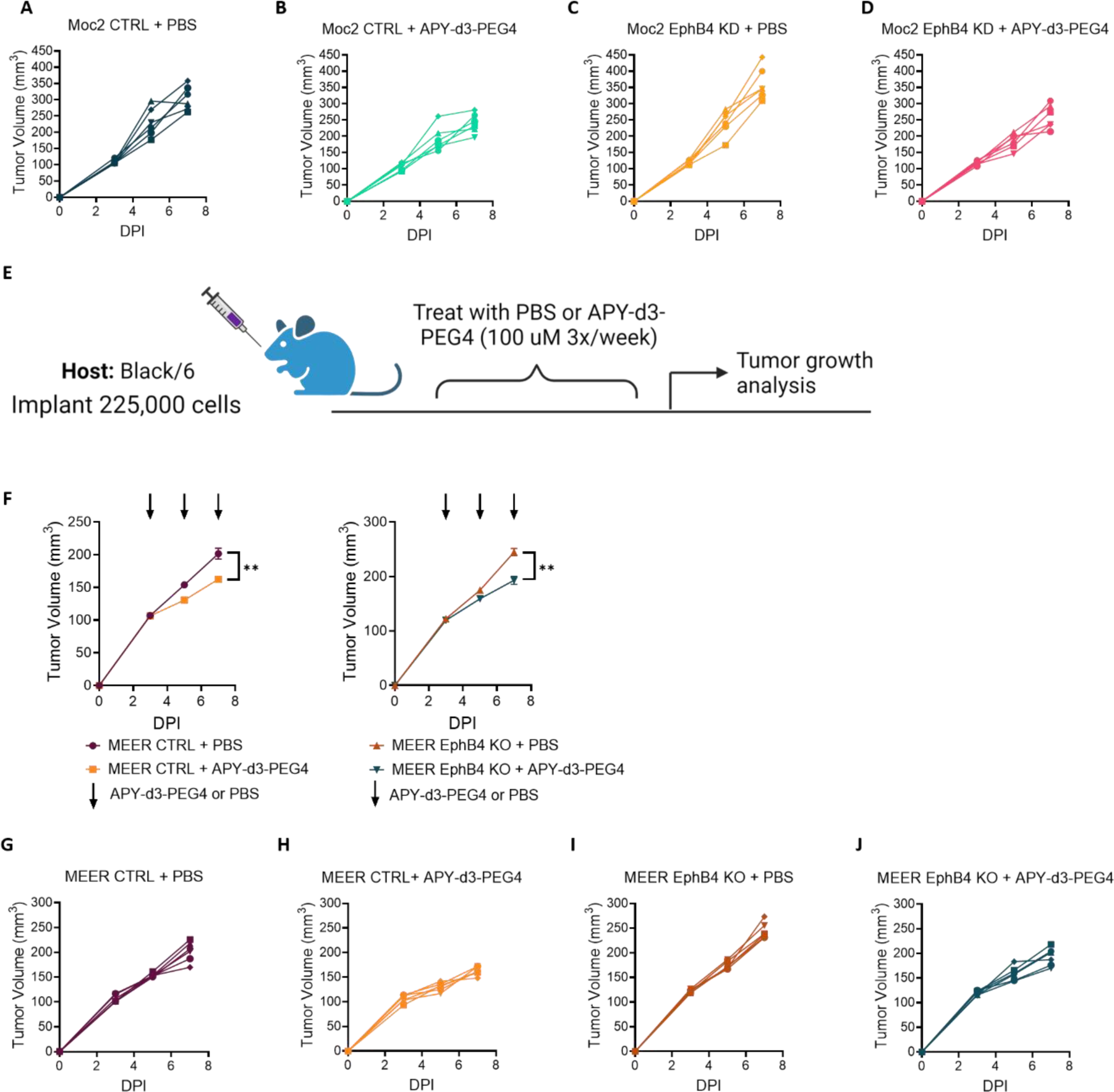

**Supplemental Figure 3.**
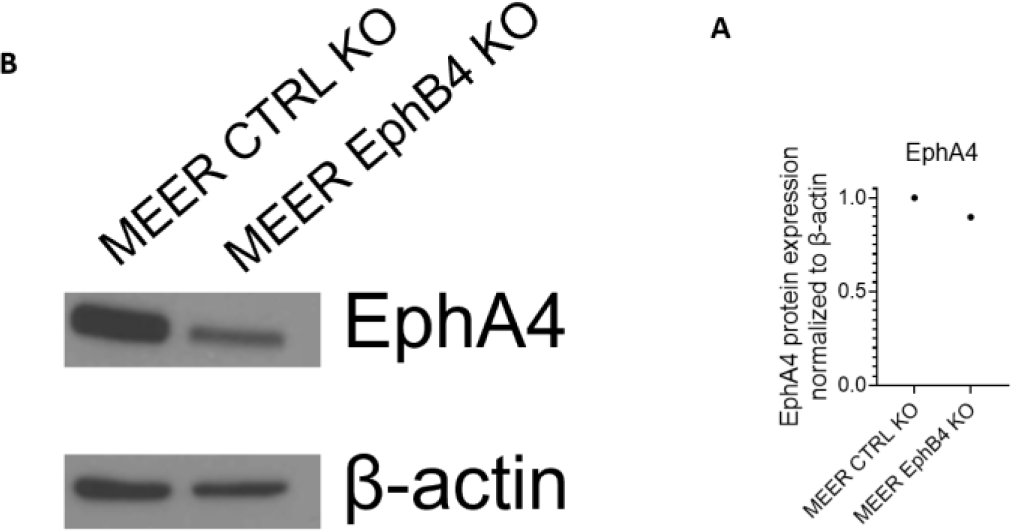

**Supplemental Figure 4.**
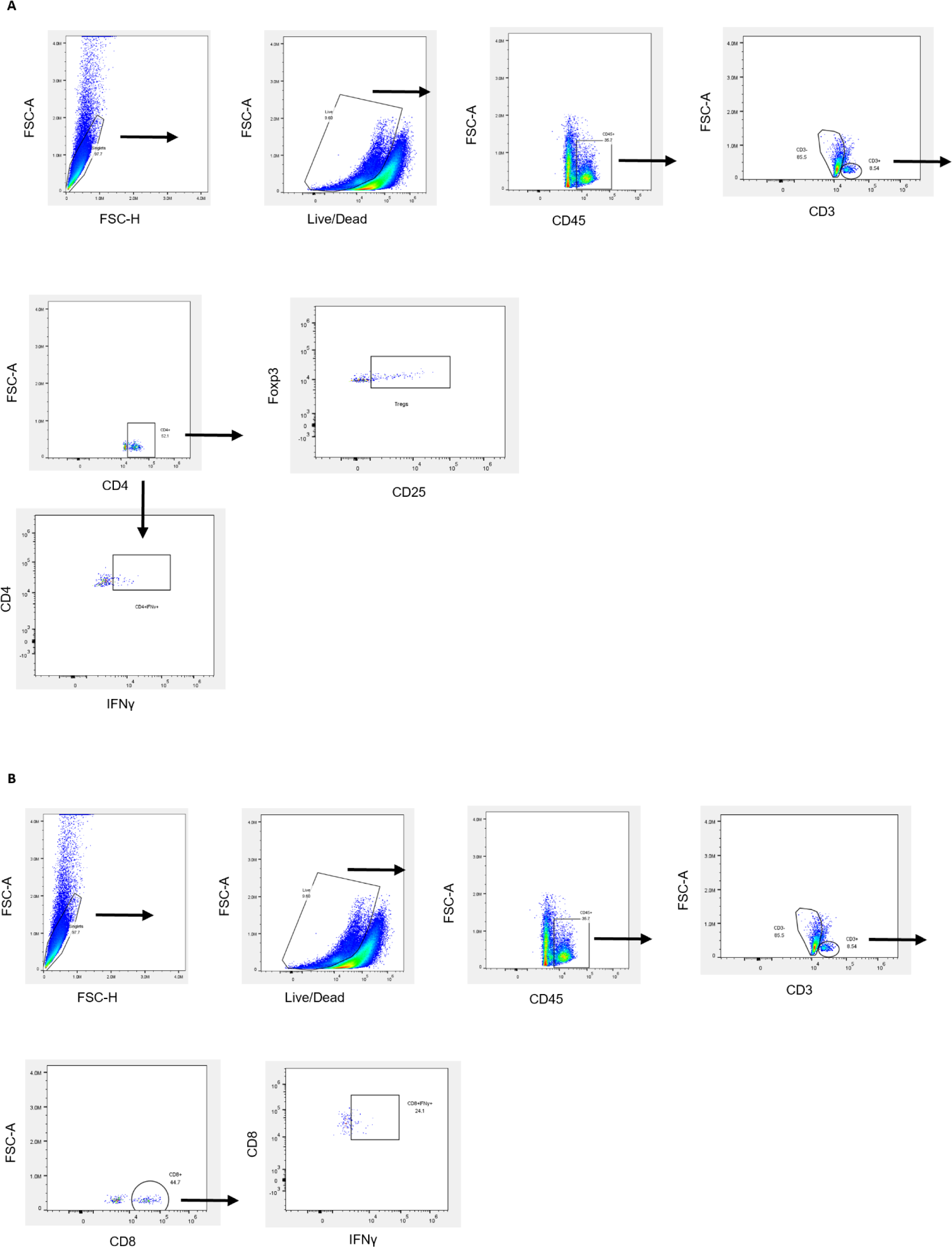

**Supplemental Figure 5.**
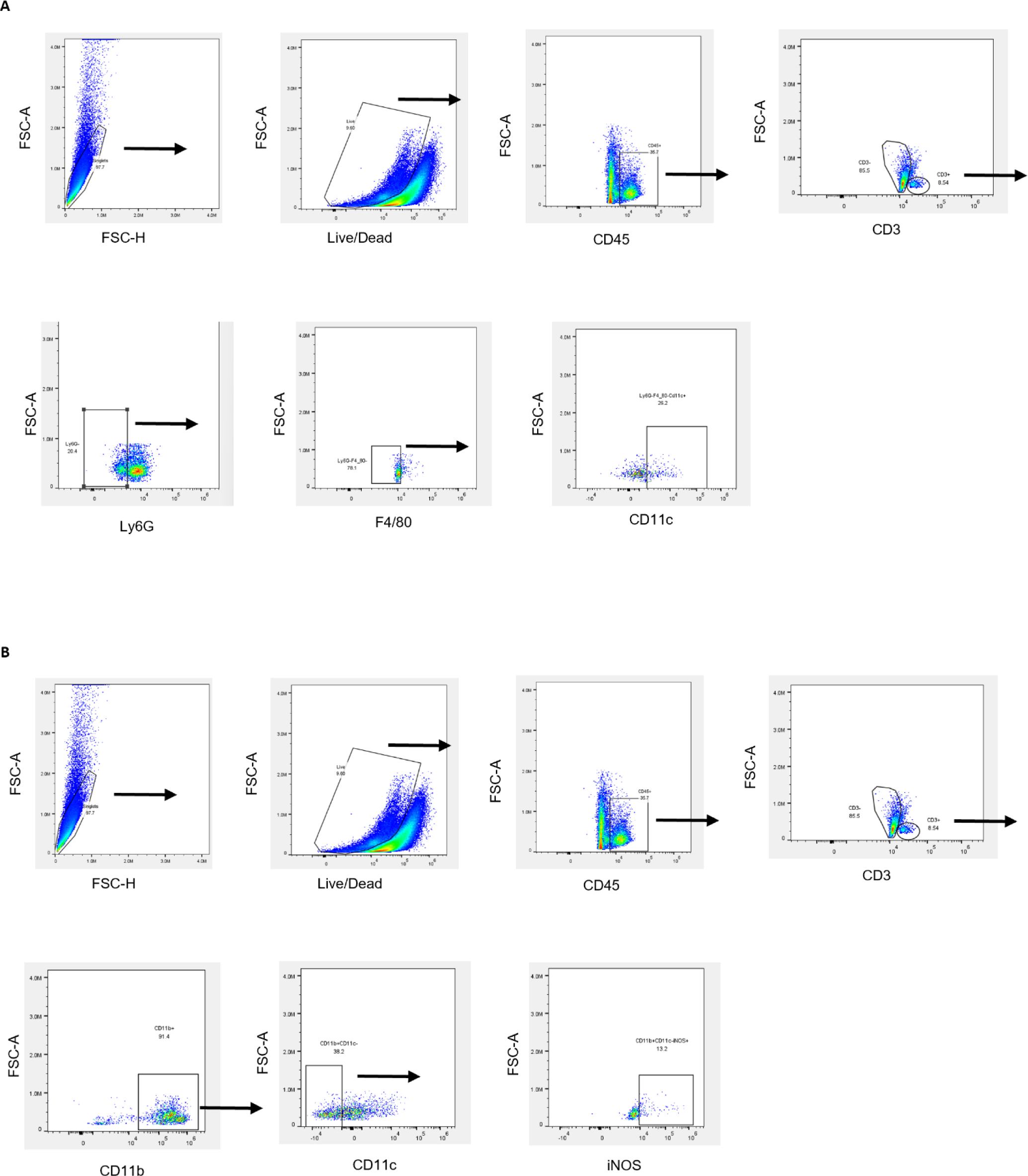

**Supplemental Figure 6.**
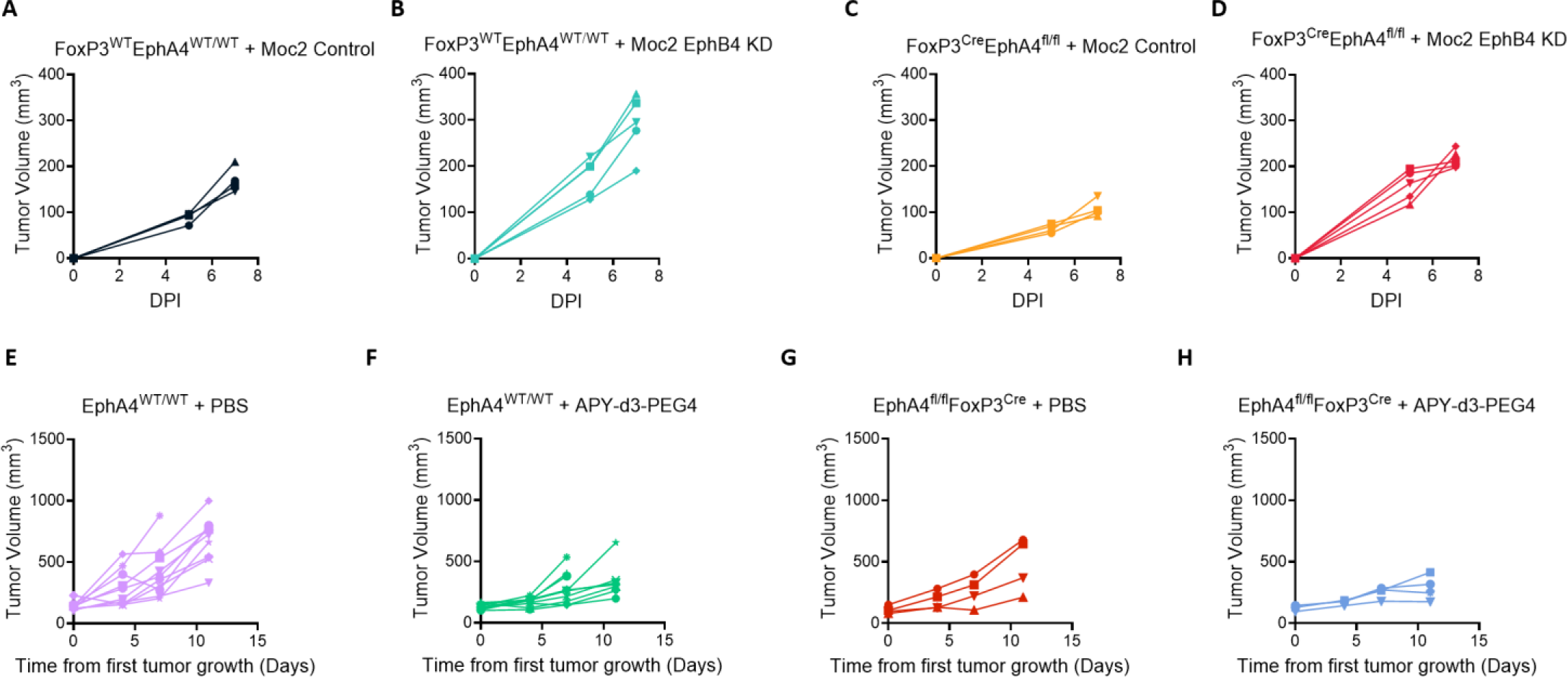

**Supplemental Figure 7.**
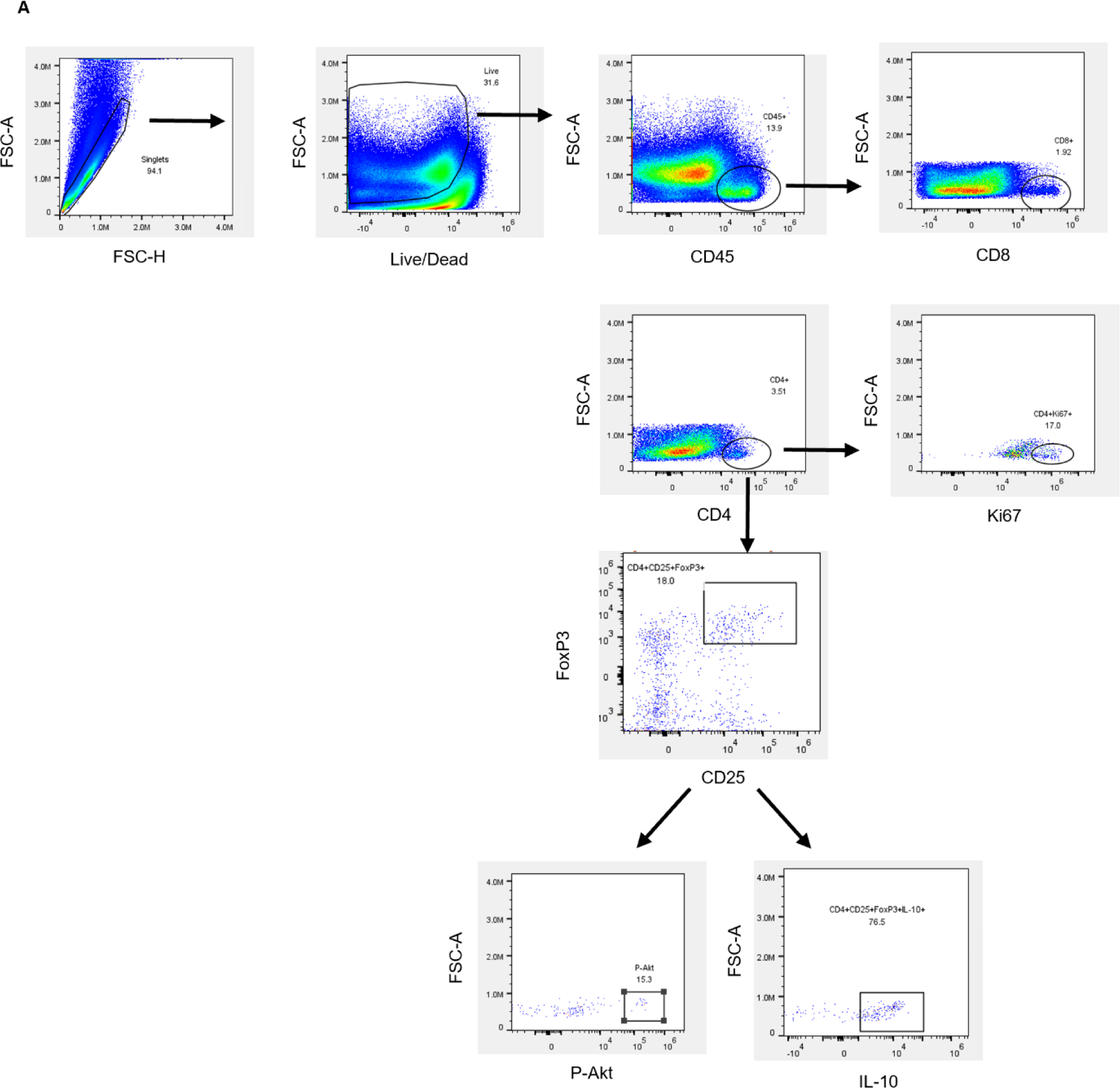

**Supplemental Figure 8.**
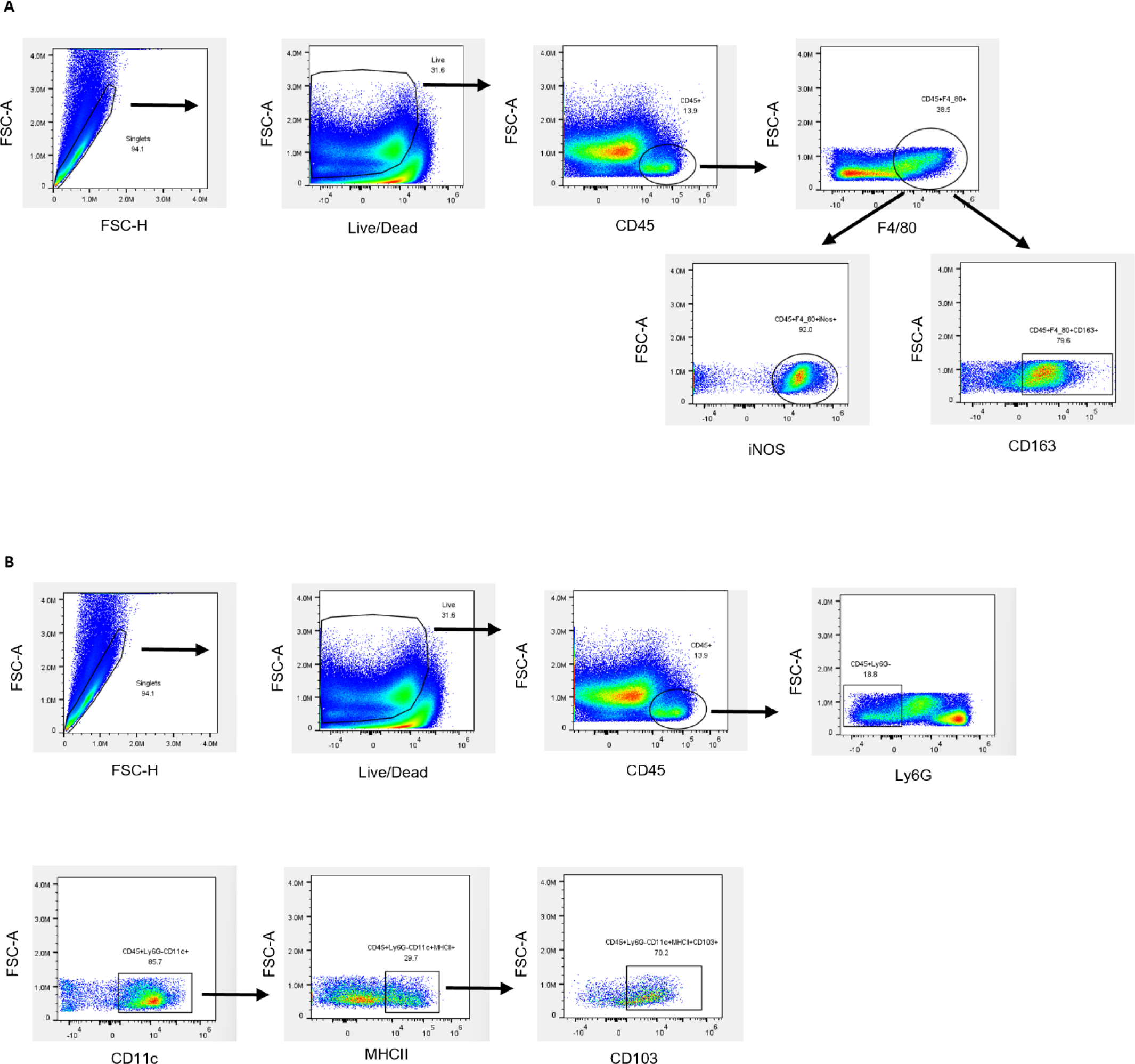

**Supplemental Figure 9.**
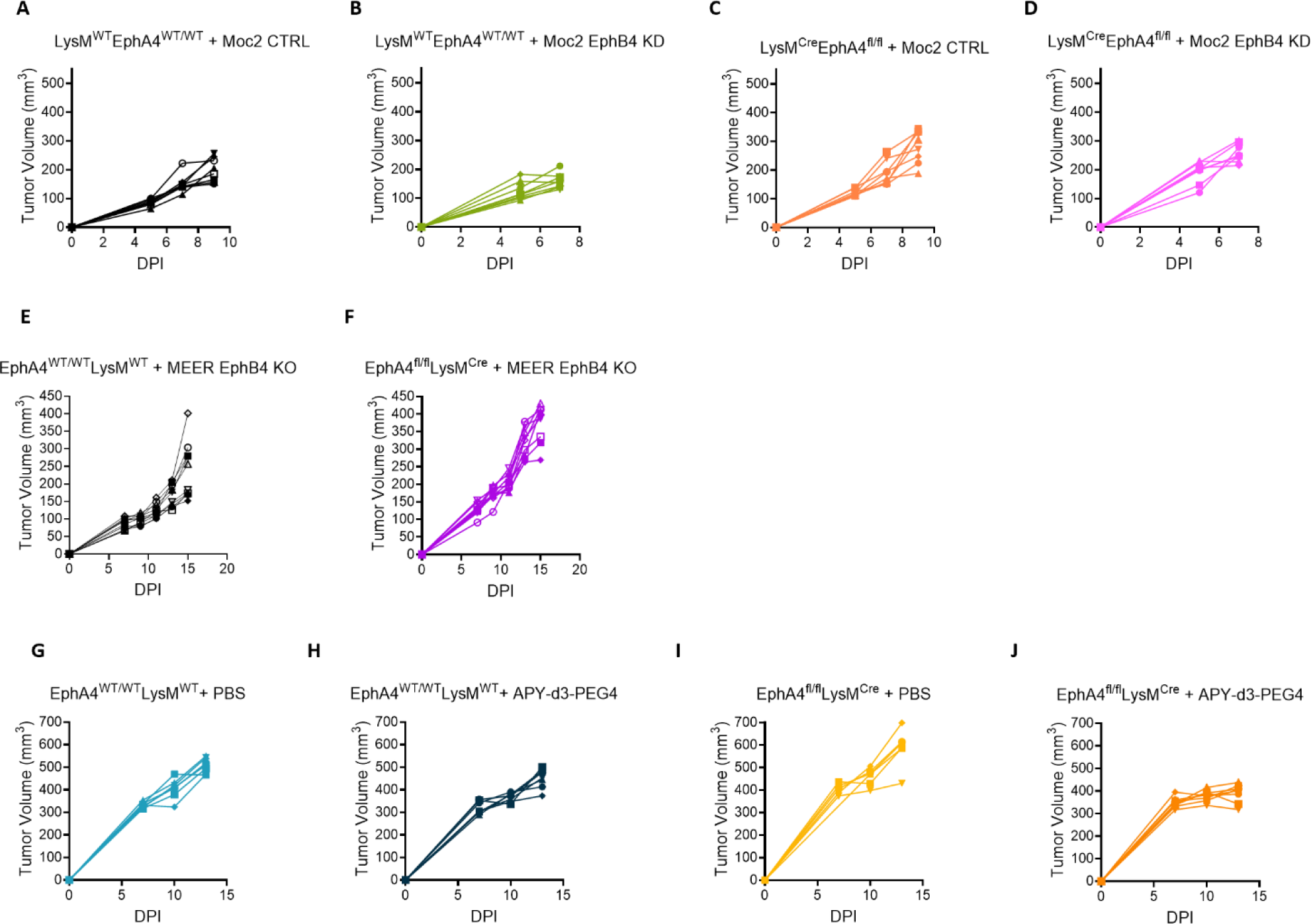

**Supplemental Figure 10.**
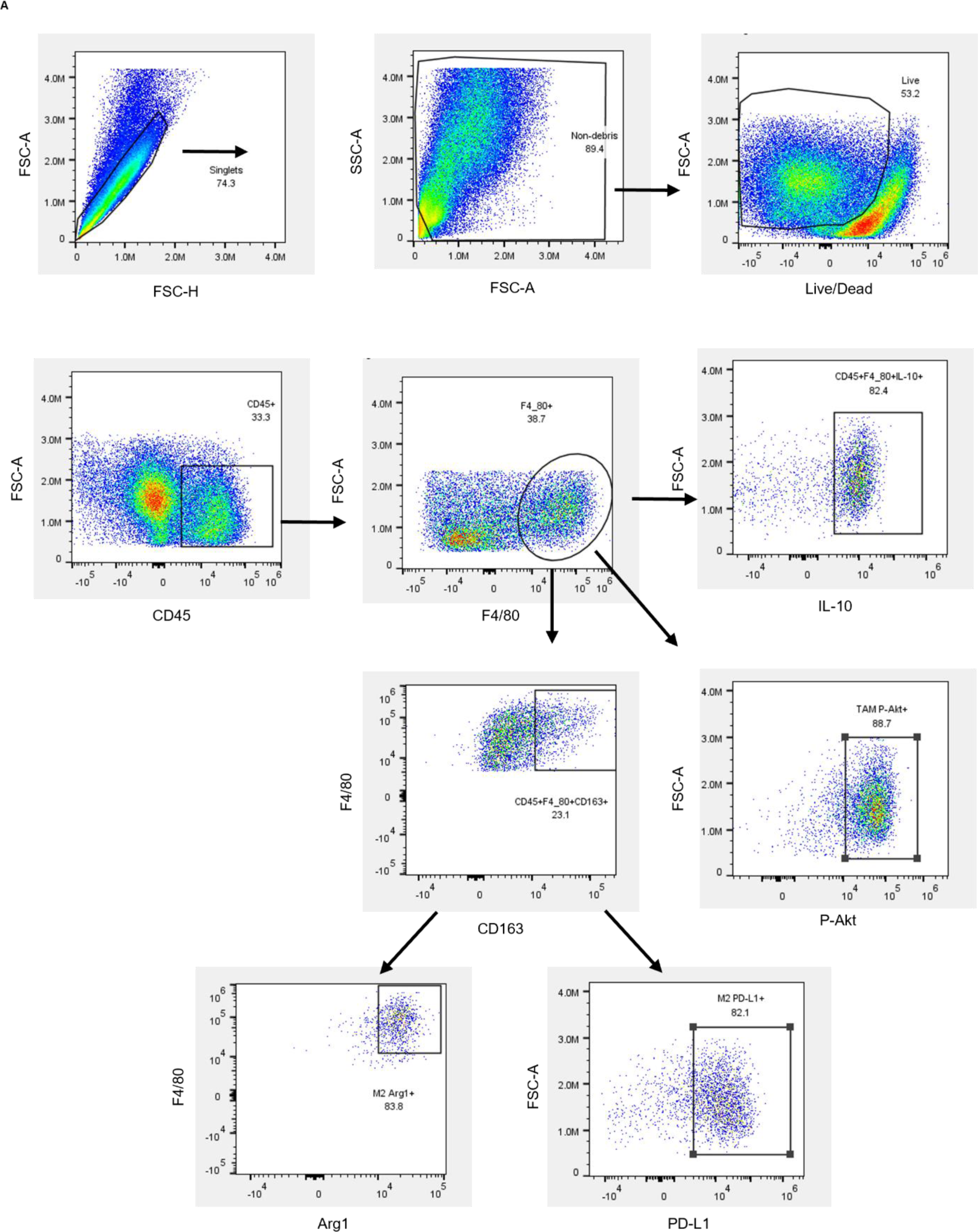

**Supplemental Figure 11.**
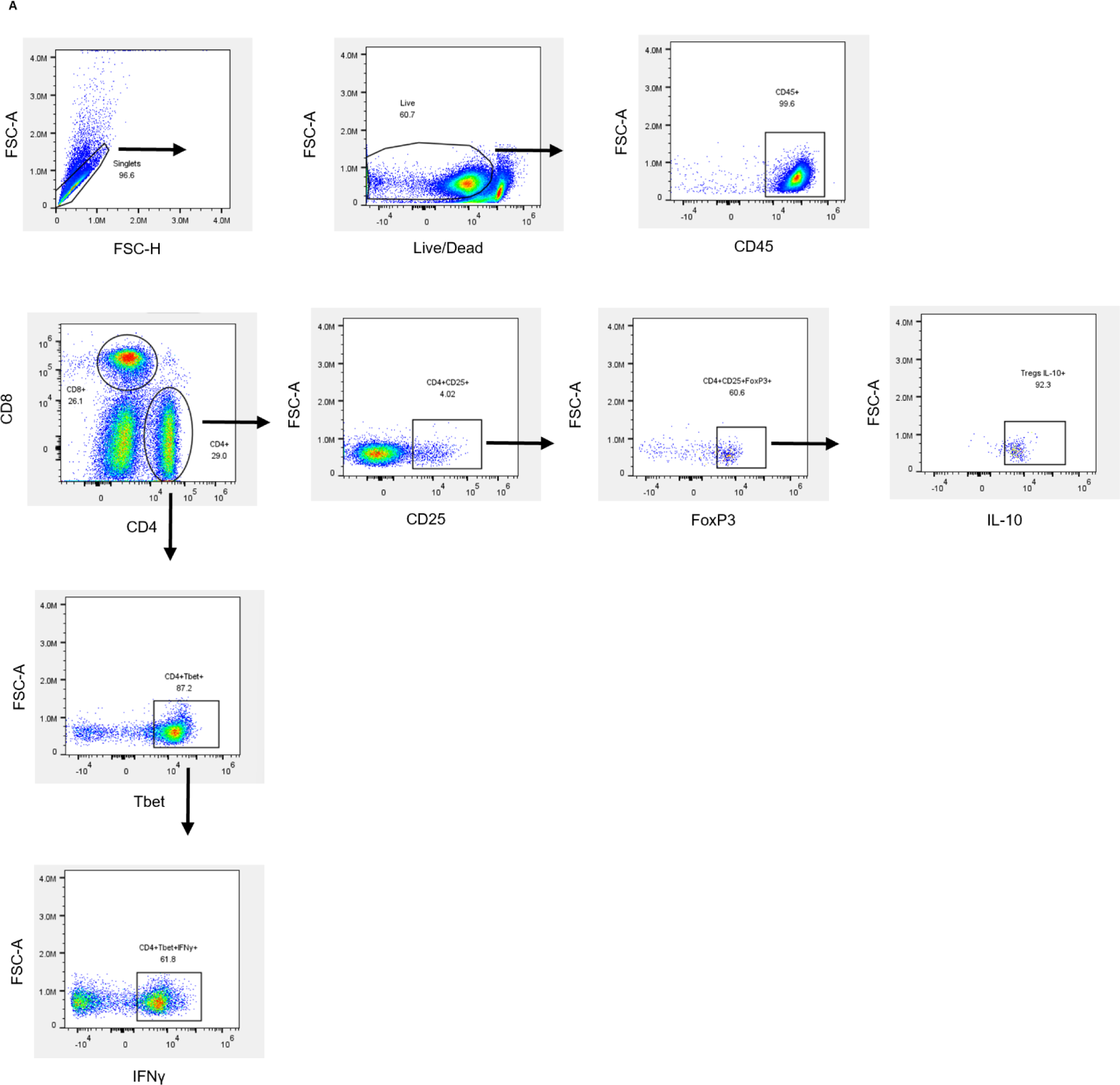

**Supplemental Figure 12.**
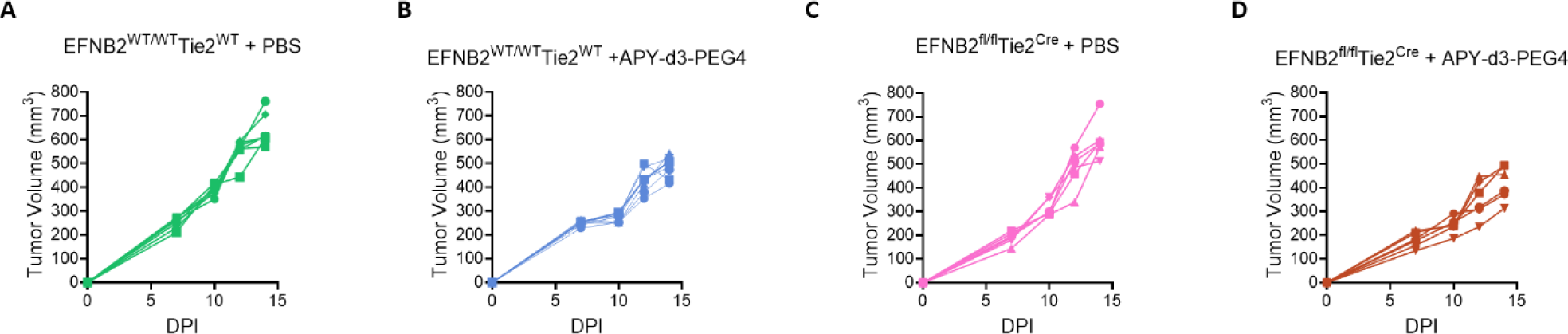

**Supplemental Figure 13.**
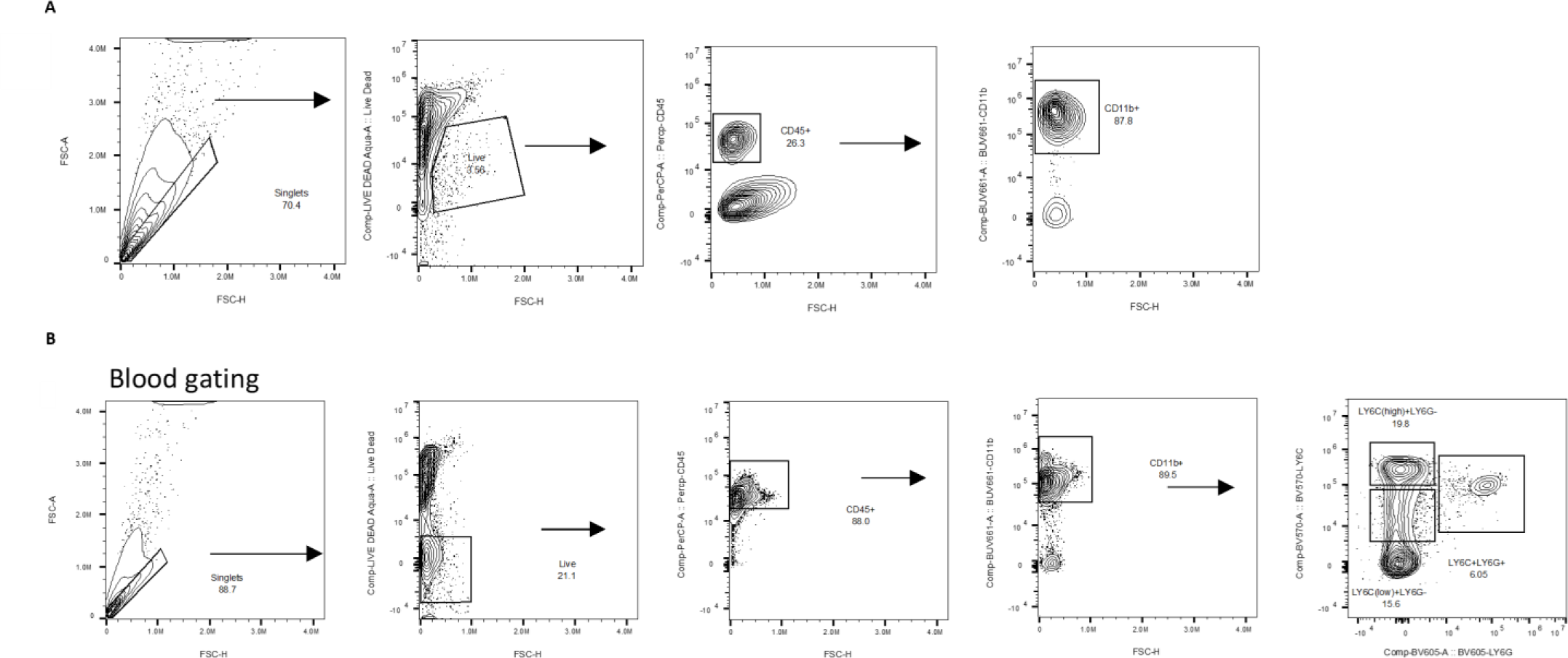

