## Supplementary material for "The pro-tumoral and anti-tumoral roles of EphA4 on T regulatory cells and tumor associated macrophages during HNSCC tumor progression": All supplemental figures will be used for the link to the file on the preprint site

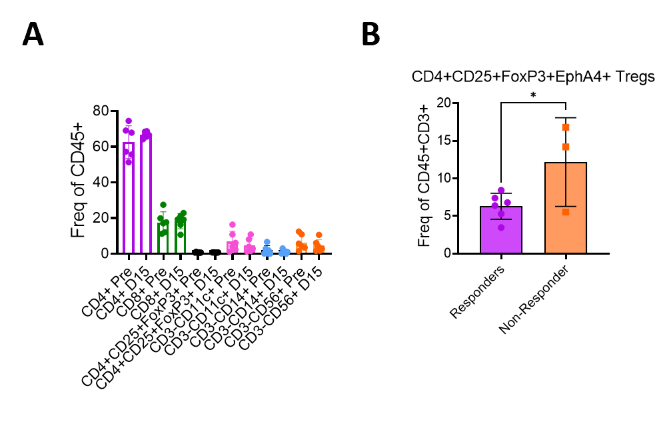

**Supplemental Figure 1. EphA4 expression in OCSCC patients. A)** CyTOF was performed on PBMCs. EphA4 expression in various leukocytes from pretreatment (Pre) and 15 days post Sitravatinib treatment (D15) time points. **B)** Blood from HPV-unrelated stage III/IV HNSCC patients in phase I/Ib clinical trial^20^ was analyzed using CyTOF. Blood was collected prior to durvalumab, surgery, and radiation therapy/chemotherapy. Orange represents patients that did not respond to combination treatment and purple represents the patients that reached complete response and major pathological response. Statistical significance was determined by Mann-Whitney T test.

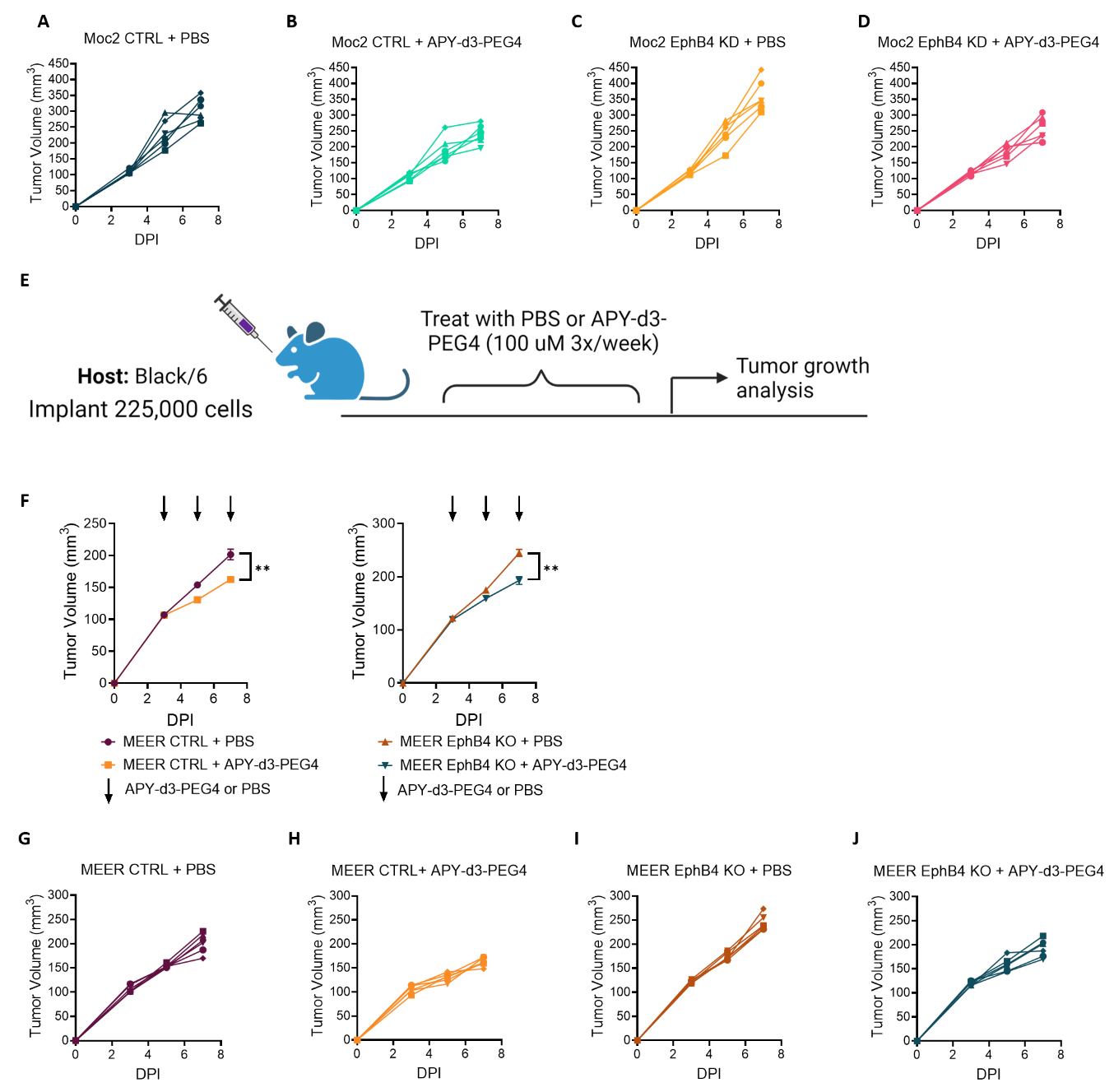

**Supplemental Figure 2. APY-d3-PEG4 limits *in vivo* tumor growth in two HNSCC orthotopic models. A-D)** Spider plots showing tumor growth for Figure 1B. Tumors were measured with calipers. **E)** Schematic of experimental design with MEER EphB4 knockout and control tumors. MEER control and MEER EphB4 knockout cells were implanted in the buccal region of C57BL/6 mice followed by treatment of PBS or 250 μL 100 μM APY-d3-PEG4 treatment three times per week by intraperitoneal injection. **F)** Tumor growth curve, caliper measurements were used to measure tumor growth. Significance was determined using a parametric two-tailed T test. **G-J)** Spider plots showing tumor growth for supplemental figure 1F.

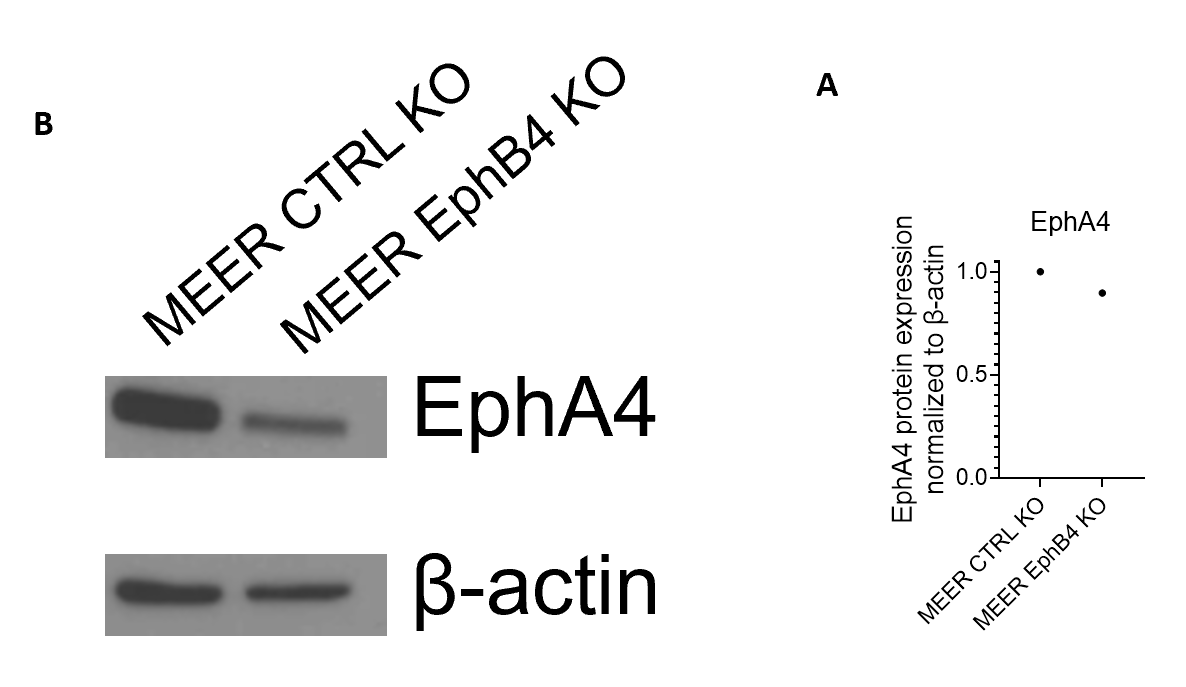

**Supplemental Figure 3. EphA4 expression does not compensate for EphB4 loss in the MEER model. A)** RNA-sequencing data on MEER control and MEER EphB4 knockout cell lines. **B)** Western blot of MEER control and MEER EphB4 knockout cell lines probed with EphA4 (120 kDa) and β-actin (42 kDa) (n = 1). **C)** Quantification of EphA4 protein expression normalized to β-actin expression (n = 1).

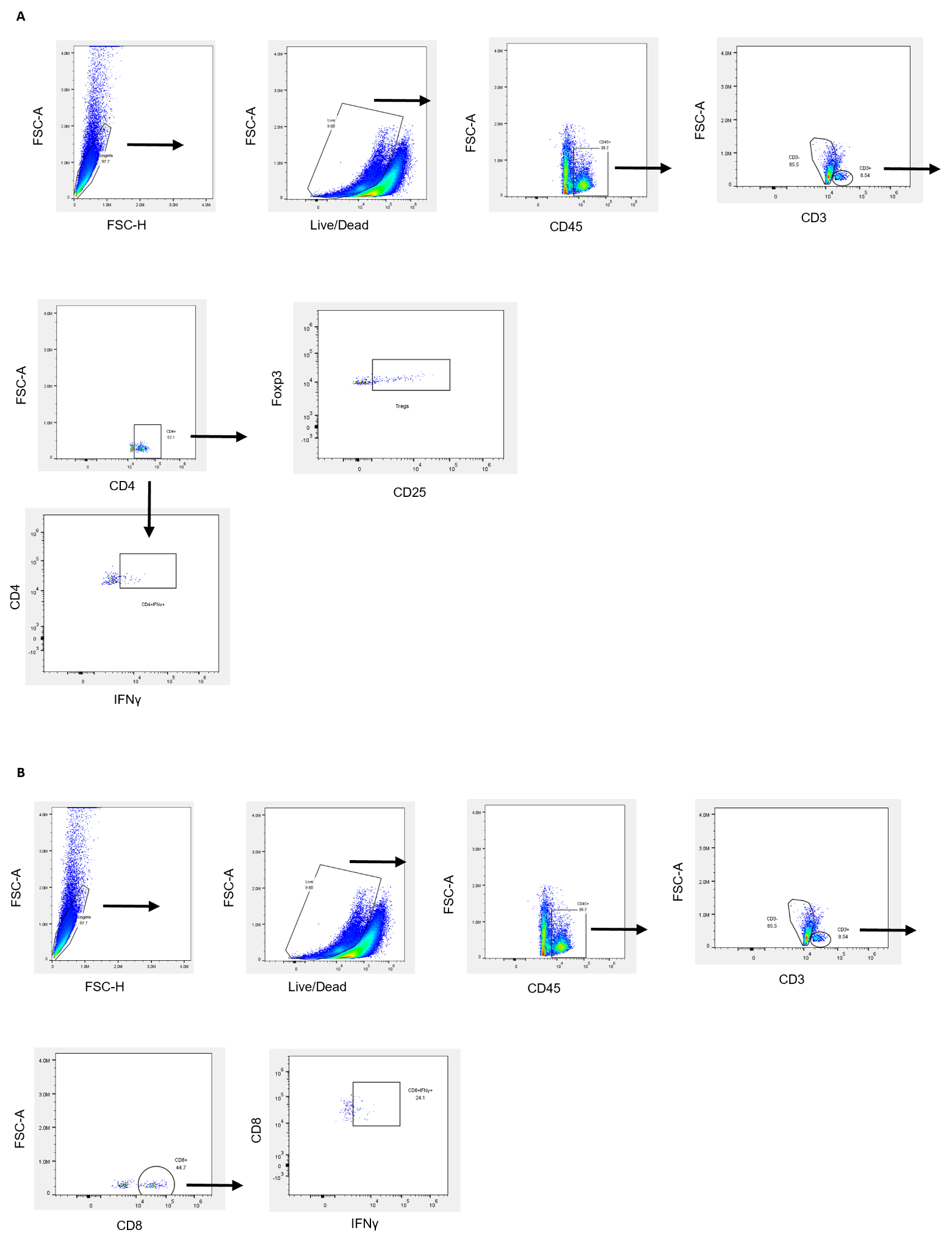
**Supplemental figure 4. Gating schematic for T cells. A)** Gating schematic for CD4+ T cells and Tregs. **B)** Gating schematic for CD8+ T cells.

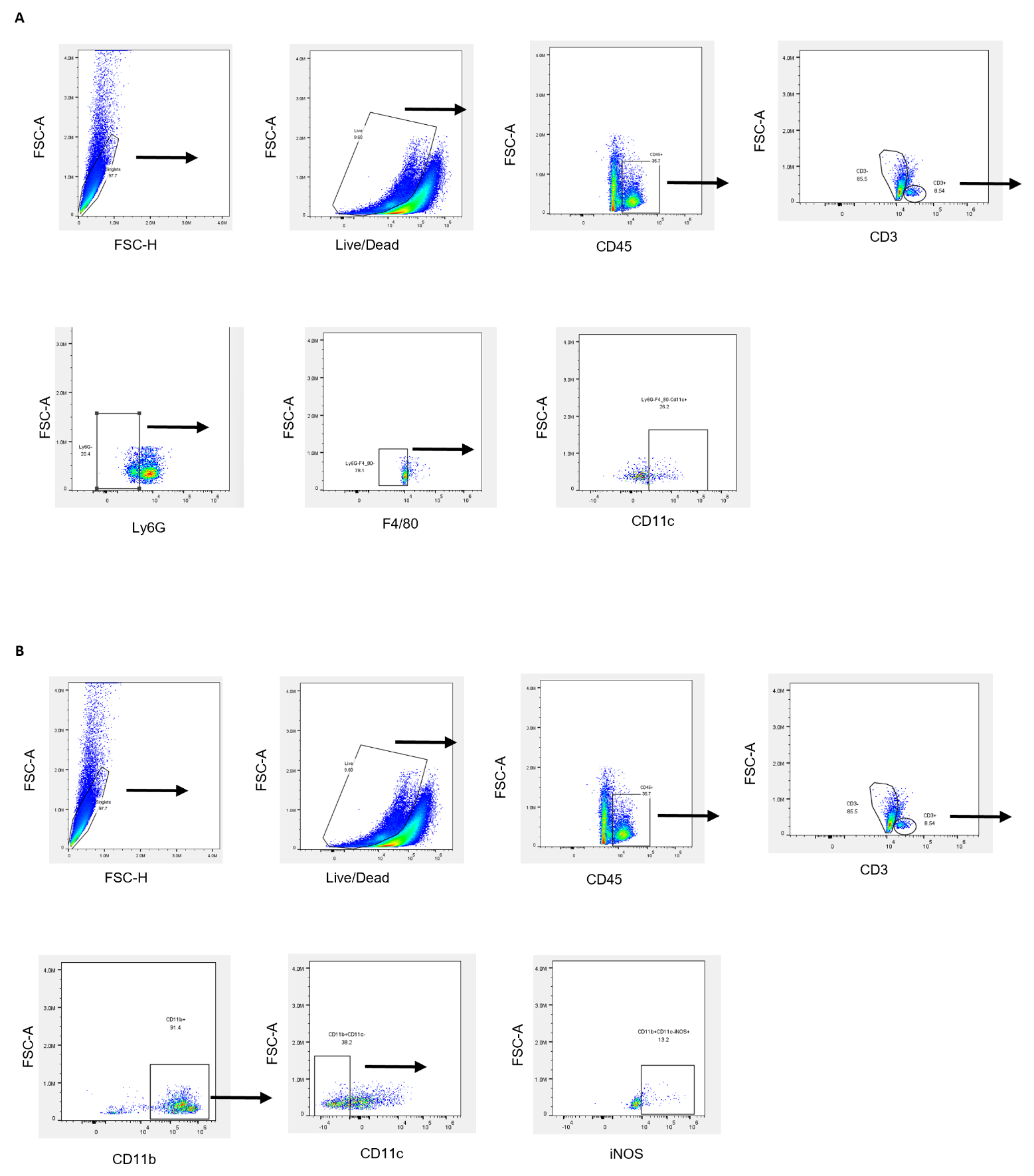

**Supplemental figure 5. Gating schematic for DCs and TAMs. A)** Gating schematic for DCs. **B)** Gating schematic for TAMs.

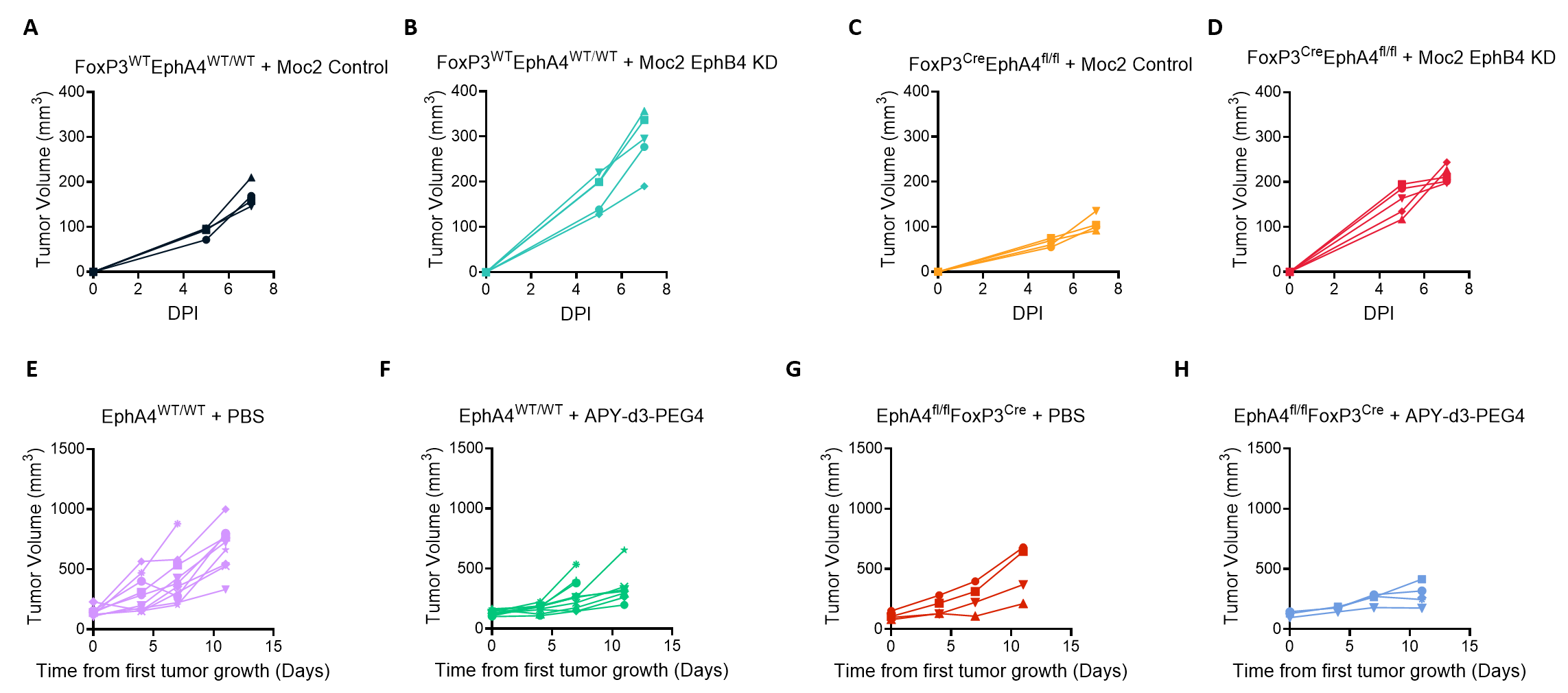

**Supplemental figure 6. EphA4 expression on Tregs mediates in part mediates accelerated tumor growth *in vivo*. A-D)** Spiders plots showing tumor growth for tumor growth curve in Figure 2b. **E-H)** Spider plots showing tumor growth for tumor growth curve in Figure 2Q.

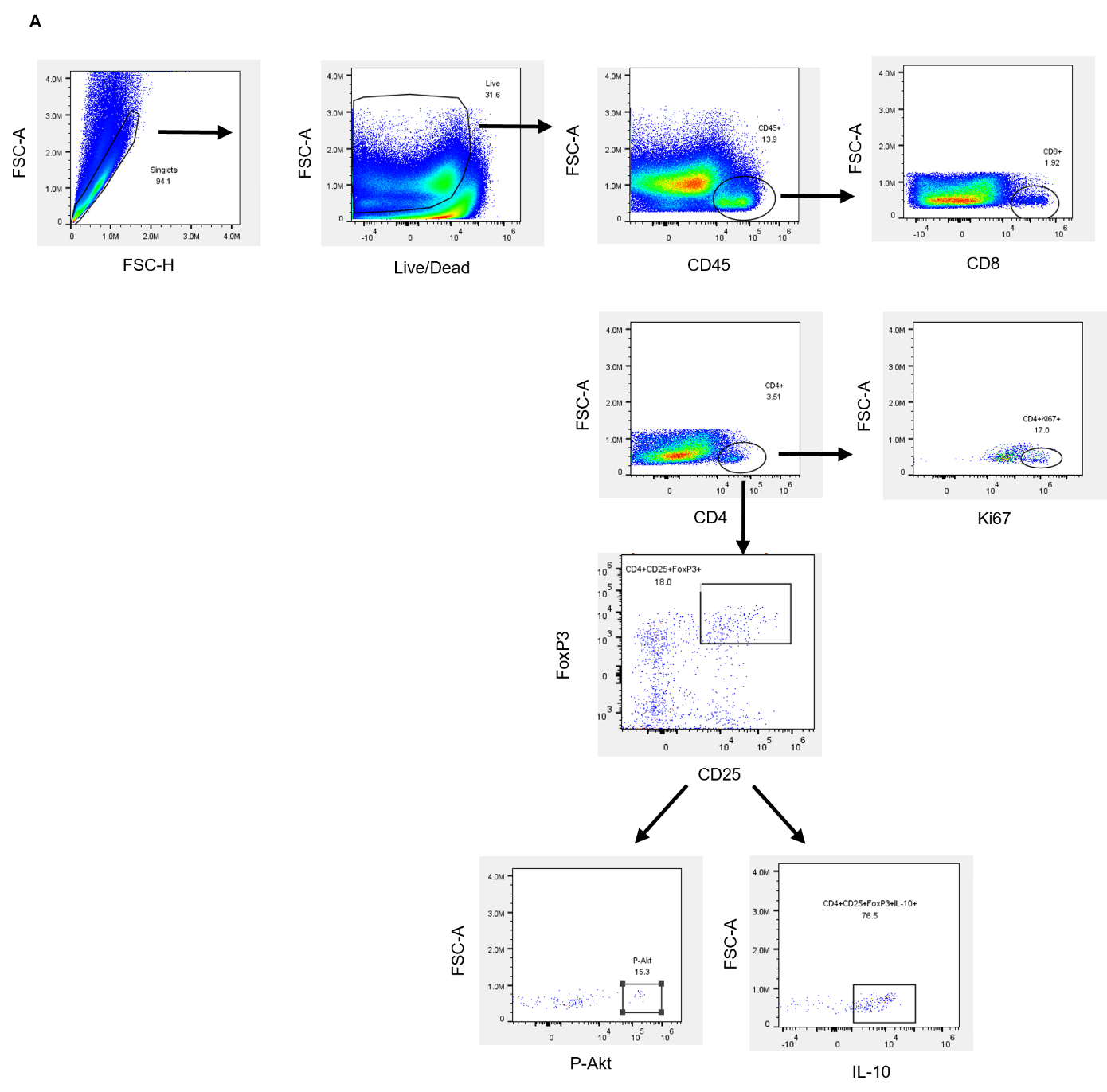

**Supplemental figure 7. Gating schematic for T cells. A)** Gating schematic for CD4+ T cells and Tregs.

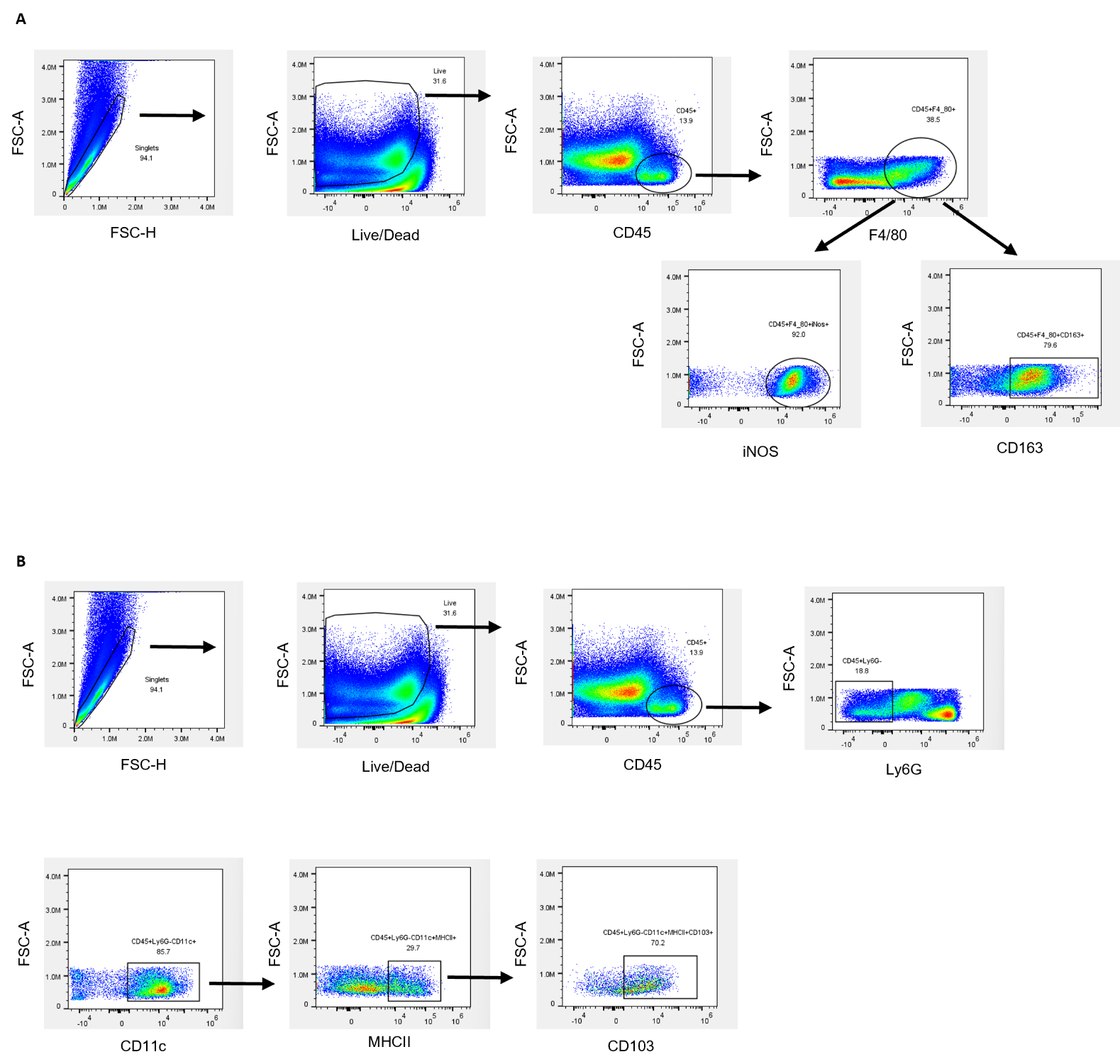

**Supplemental figure 8. Gating schematic for TAMs and DCs. A)** Gating schematic for TAMs. **B)** Gating schematic for DCs.

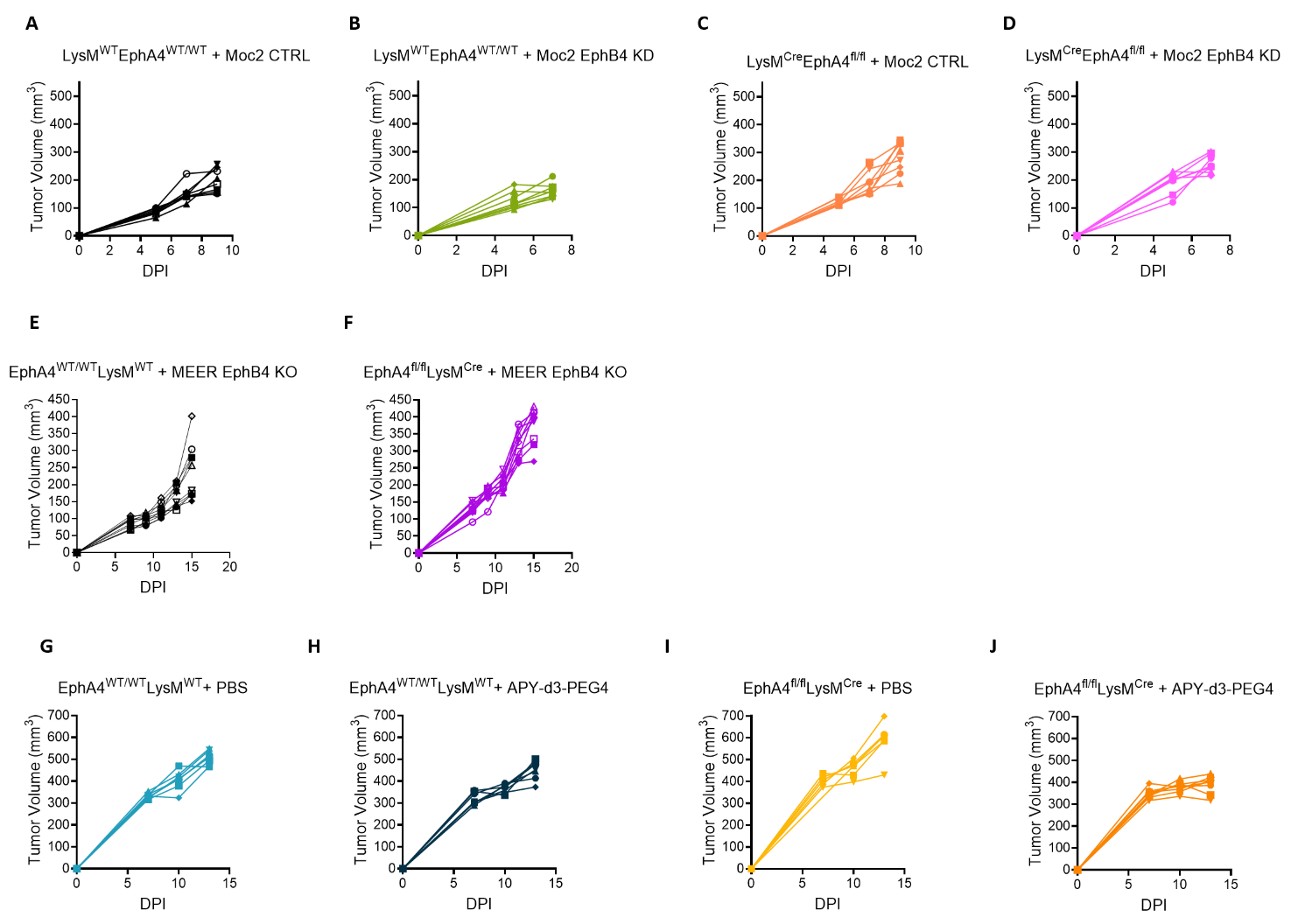

**Supplemental Figure 9. Loss of EphA4 on tumor associated macrophages promotes accelerated tumor growth. A-D)** Spiders plots showing tumor growth for tumor growth curve in Figure 3B. E-F) Spiders plots showing tumor growth for tumor growth curve in Figure 3D. **G-J)** Spiders plots showing tumor growth for tumor growth curve in Figure 3S.

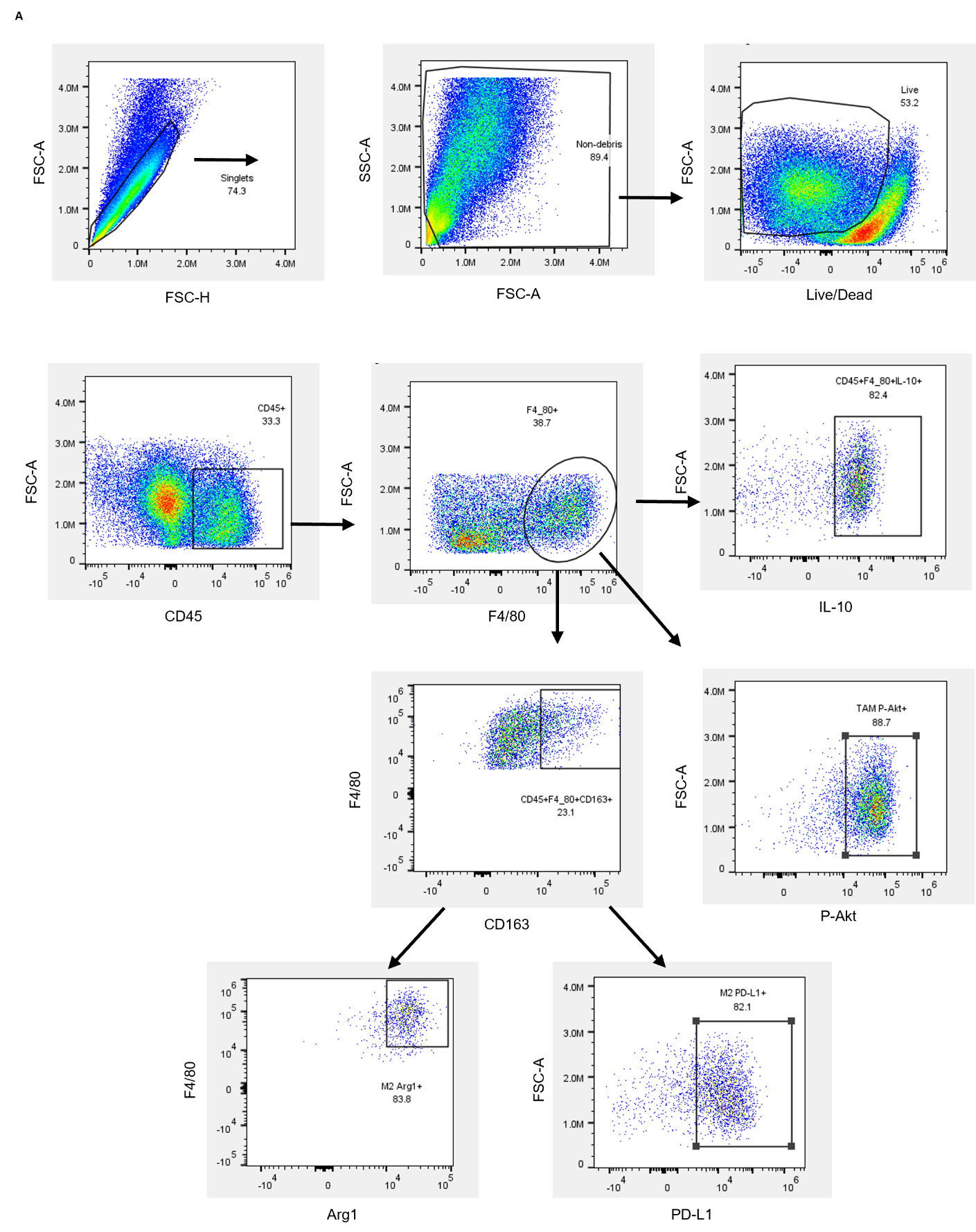

**Supplemental figure 10. Gating schematic for TAMs. A)** Gating schematic for TAMs.

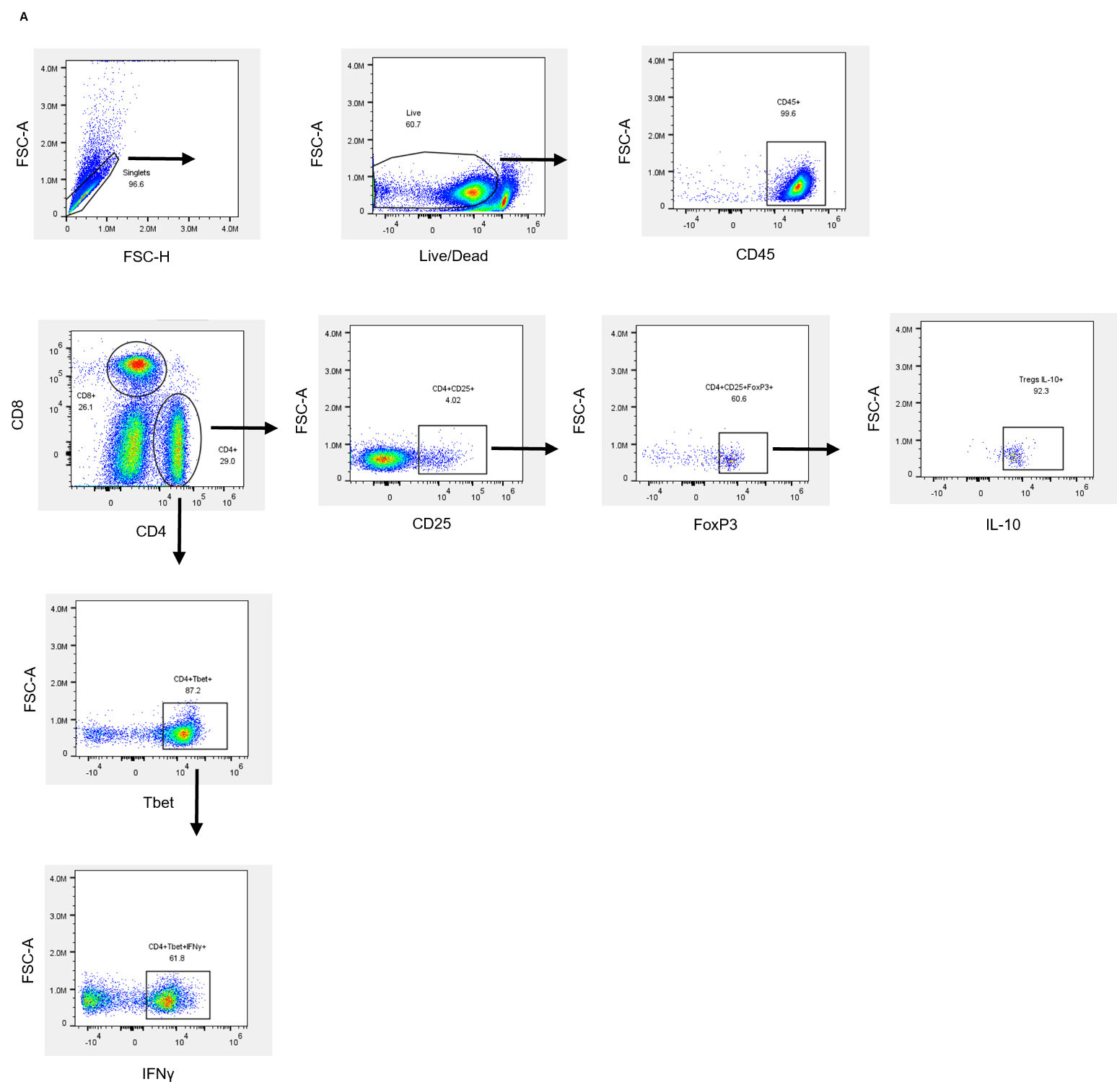

**Supplemental figure 11. Gating schematic for T cells. A)** Gating schematic for CD4+ T cells and Tregs.

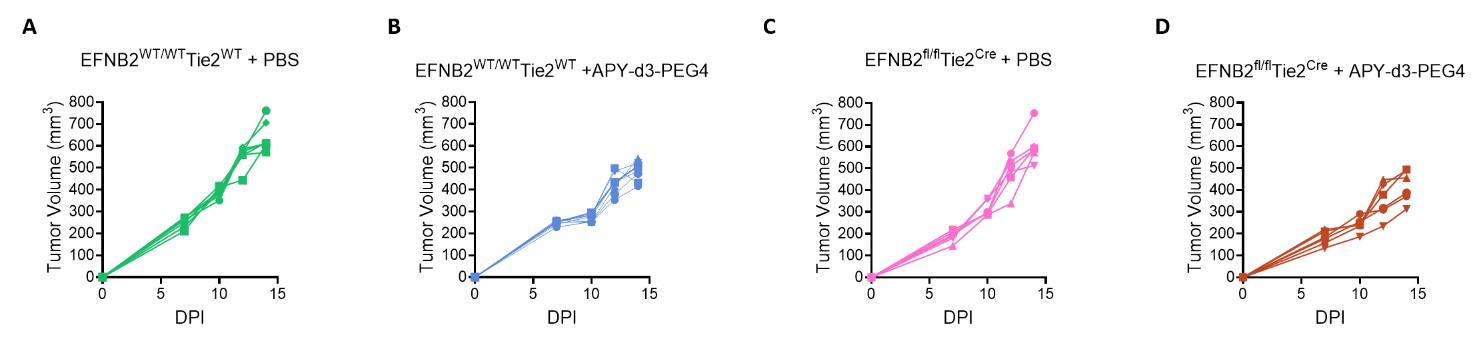

**Supplemental Figure 12. Dual inhibition of EphA4 and ephrinB2 leads to reduced tumor growth. A-D)** Spiders plots showing tumor growth for tumor growth curve in Figure 4K.

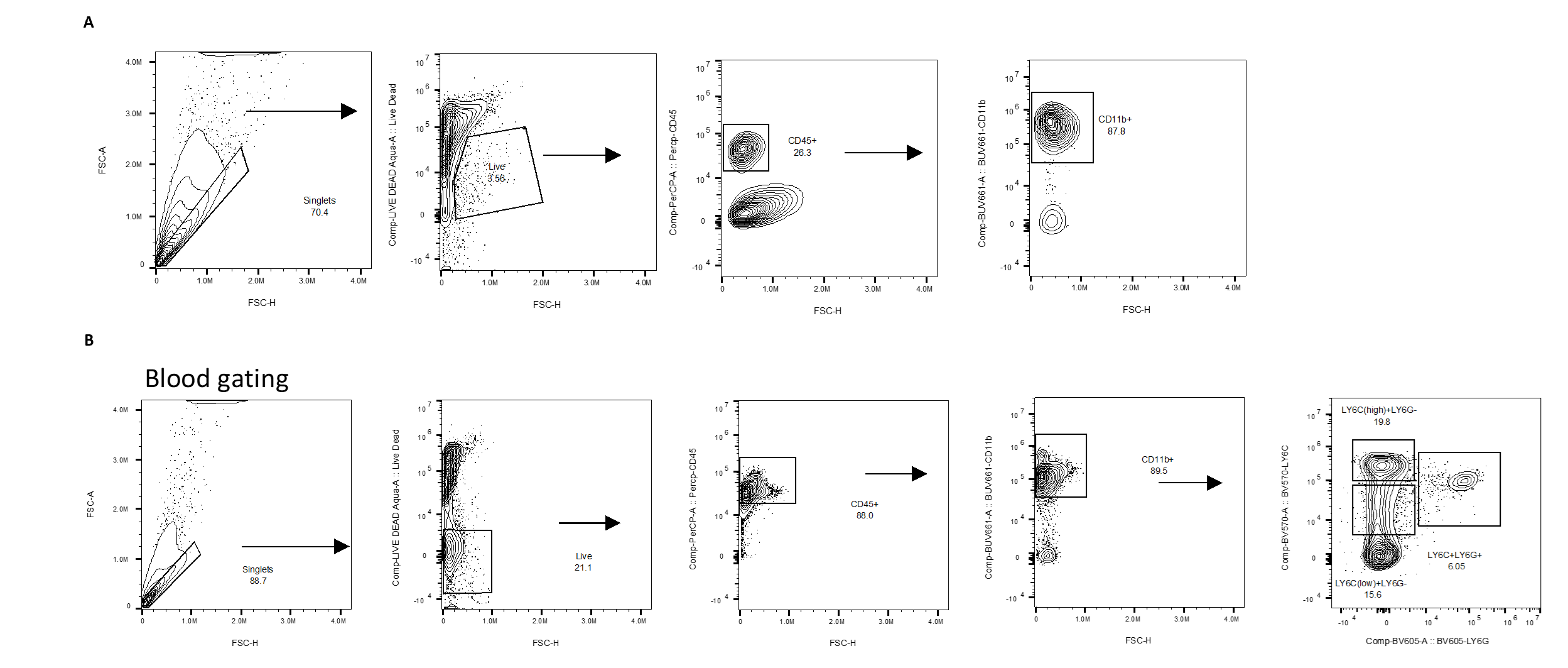

**Supplemental figure 13. Gating schematic for TAMs and monocytes. A)** Gating schematic for TAMs. **B)** Gating schematic for monocytes.
